# Differential expression of miRNAs between Young-Onset and Late-Onset Indian colorectal carcinoma patients

**DOI:** 10.1101/2023.07.19.549649

**Authors:** Sumaiya Moiz, Varsha Mondal, Debarati Bishnu, Biswajit Das, Bodhisattva Bose, Soumen Das, Nirmalya Banerjee, Amitava Dutta, Krishti Chatterjee, Srikanta Goswami, Soma Mukhopadhyay, Sudarshana Basu

## Abstract

Early-Onset Colorectal Carcinoma (EOCRC) is a growing concern as reports indicate a worldwide increase in the incidence of CRC among young adults (<50 years old). In an effort to understand the different mode of pathogenesis in young-onset CRC, we performed a pilot study wherein we looked at colorectal tumors from both young (< 50 years old) and old patients (>55 years old) and screened them to eliminate tumors positive for Microsatellite Instability (MSI) and showing activation of the Wnt pathway, known canonical factors in CRC pathogenesis. RNA isolated from EOCRC and Late-Onset (LOCRC) tumors and paired normal tissues without MSI, nuclear β-catenin and APC mutations were sent for small RNA seq to identify miRNA alterations between the two subsets. Comparative analysis revealed differential expression of 23 miRNAs specific to EOCRC and 11 miRNAs specific to LOCRC. We validated the top 10 EOCRC DEMs in TCGA-COAD cohorts followed by validation in additional EOCRC and LOCRC cohorts. Our integrated analysis revealed upregulation of hsa-miR-1247-3p, hsa-miR-148a-3p and hsa-miR-27a-5p and downregulation of hsa-miR-326 between the two subsets. Experimentally validated targets of the above miRNAs were compared with differentially expressed genes in the TCGA dataset to identify targets with physiological significance in EOCRC development. Our analysis revealed downregulation of epithelial gene expression and intercellular junction proteins potentially leading to dissolution of epithelial intercellular junctions in EOCRC development and EMT progression. Upregulated targets included genes whose expression have been reported to correlate with CRC tumor invasion, liver metastasis, disease recurrence and poor prognosis.

## Introduction

Colorectal cancer (CRC) accounted for 10% of all new cancer cases in 2020 and is the third-most prevalent cancer in the world (https://gco.iarc.fr/today/data/factsheets/populations/900-world-fact-sheets.pdf). As per data from Globocan 2020, there were ∼65,358 new CRC cases in 2020 in India making it the fifth most prevalent cancer in the country. Projections taking into account aging, population growth and human development estimate that by 2040, the incidence of colorectal cancer in India will increase by more than 60% (Globocan 2020). The number of young people (0-49 years) expected to be diagnosed with CRC has been estimated to increase by more than 8% between 2020 and 2025.

Age represents the primary risk factor for CRC, although the cumulative risk for early-onset CRC (0-49 years old) in India has increased (for males) from 0.06 in 1986 to 0.13 in 2012. Analysis of CRC incidence via the Surveillance, Epidemiology, and End Results (SEER) data from 2000 to 2019 revealed that adults aged less than 50 years were noted to have an average of 2.4% annual increase in CRC incidence rates, while adults above the age of 65 years had an average of −3.4% change in CRC incidence. Thus, although there is a general decrease in the overall incidence of colorectal cancer, the incidence of the disease in young adults (<50 years) has increased worldwide. This has prompted the ACS (American Cancer Society) to revise the standard age for CRC risk screening to 45 years from 50 years (**1**).

Consensus exists that Early Onset CRC (EOCRC) is pathologically, anatomically, metabolically and biologically different from Late Onset CRC (LOCRC) and hence should be investigated and managed differently (**2**, **3**, **4**). EOCRC tumours were found to be mostly located in the distal colon (80%), particularly the sigmoid colon and the rectum, with a higher prevalence of adverse histological factors such as signet ring cell differentiation, venous invasion, and perineural invasion (**5**). The tumours lacked frequent activating BRAF or KRAS mutations, suggesting that the molecular events in tumour development differed with respect to late-onset group. Additionally, EOCRC was not frequently associated with precursor adenomatous lesions suggesting that the classical adenoma to carcinoma pathway of molecular events (**3**, **6**) does not occur in this patient subset.

Recently gene expression analyses have identified putative targets that indicate altered pathways in EOCRC. The MAPK signalling pathway appeared to be deregulated in the early-onset sporadic group as compared to PI3K-AKT in the late-onset group (**7**). Bioinformatics analysis on microarray data sets identified strong implication of molecular mechanisms involved in vascular smooth muscle contraction signaling pathway (**8**). Early onset sporadic tumours lacking canonical genetic aberrations like MSI and WNT-β-catenin activation were found to be enriched in Ca2+/NFAT pathways (**9**, **10**). High throughput RNA sequencing of EOCRC tumours followed by validation by RT-qPCR identified significant upregulation of TNS1 and MET (**11**). However, miRNAs deregulated in EOCRC patients have not yet been explored, especially in Indian patients. miRNAs are important regulator molecules that may act as either tumor suppressor or oncogenes depending on the cellular environment in which they are expressed. Analysis of miRNA expression profiles and their predicted target genes can indicate the aberrant physiology of a system and may be targeted in therapy or used as biomarkers for diagnostic purposes.

We performed a pilot study wherein we looked at the overall miRNA profile of Indian CRC tumours which had been screened to exclude hereditary CRC syndromes like Lynch Syndrome and Familial Adenomatous Polyposis. Microsatellite Stable (MSS) tumours without nuclear β-catenin and APC mutations (mutations in the mutation cluster region (MCR) from codon 1,250 to 1,464) were divided into two subsets: EOCRC (disease onset <50years) and LOCRC (disease onset >55 years). We investigated miRNA alterations between our EOCRC and LOCRC subsets by genome-wide small RNA sequencing. Differentially expressed miRNAs (DEMs) were validated by quantitative real-time PCR in additional EOCRC and LOCRC patient cohorts followed by subsequent bioinformatic analysis of the validated miRNAs to identify novel deregulated pathways in EOCRC. Our analysis outcome identifies miRNAs and pathways differentially expressed between EOCRC and LOCRC. To the best of our knowledge, this study is the first to compare miRNA expression between EOCRC and LOCRC patients in India.

## MATERIALS AND METHODS

### Patient recruitment

Colorectal tumour samples from histologically proven CRC patients were collected in collaboration with doctors and pathologists from the Surgical Oncology Department of Netaji Subhas Chandra Bose Cancer Hospital, Kolkata (discovery cohort) and the Departments of Surgical, Medical & Radiation Oncology, Surgical Gastroenterology and General Surgery, All India Institute of Medical Sciences (AIIMS), Rishikesh (validation cohort). All patients with histopathologically proven colorectal cancers undergoing treatment from May 2020 to May 2022 who fulfilled the inclusion and exclusion criteria were included. All necessary IEC permissions were obtained prior to sample collection. Clinicopathological information like age, gender, site, stage, and differentiation of tumour, familial history of CRC and presence of any other inflammatory bowel disease were also collected. Inclusion criteria: Patients admitted for surgical resection with biopsy proven colorectal adenocarcinoma, age upto 80 years, willing to provide written informed consent. Exclusion criteria: Patients with Familial Colorectal Carcinoma, Unable/ unwilling to give consent, Patients with cancers other than CRC, patients receiving neoadjuvant chemotherapy and/or radiotherapy.

### Biospecimen collection

Tumour and Normal tissue samples were collected in RNALater (Invitrogen) and 10% neutral buffered formalin. Samples collected in RNA Later (Invitrogen) for Nucleic acid extraction were stored at −80°C for processing at a later date. Samples stored in neutral buffered formalin were processed into FFPE blocks, sectioned into 5µm sections and adhered onto positively charged slides. Histological sections were stained with haematoxylin and eosin. All specimens with histopathological features suggestive of an inflammatory colorectal disease were excluded from the study. Reporting was done by trained histopathologists. Grossing and reporting of colectomy specimens suspicious of colorectal carcinoma were conducted according to CAP (College of American Pathologists).

### Immunohistochemistry

IHC for MMR proteins and nuclear β-catenin were done as per standard protocol. Retrieval of antigen was done in Tris-EDTA buffer (pH 9) for MMR proteins and in 10mM Sodium citrate buffer (pH 6) for nuclear β-catenin in the microwave. Slides were analysed by a trained histopathologist for detection of MSI (as per College of American Pathologists (CAP) guidelines) and nuclear β-catenin. Normal colorectal tissue was taken as internal control. Known case of MSI CRC was used as positive control for MSI detection. External control for nuclear β-catenin consisted of histologically diagnosed section of fibromatosis (Desmoid tumour). No antibody controls were taken as negative controls. The scoring of Wnt positive nuclear β-catenin expression (Wnt+) was performed according to Raman et al. (**9**). A sample was scored as Wnt positive (Wnt+) if β-Catenin nuclear stain was observed in more than 35% tumor epithelial cells and Wnt negative (Wnt−) if nuclear stain was detected in less than 25% cells. IHC images were captured using the Olympus BX53 biological microscope.

### List of primary antibodies used

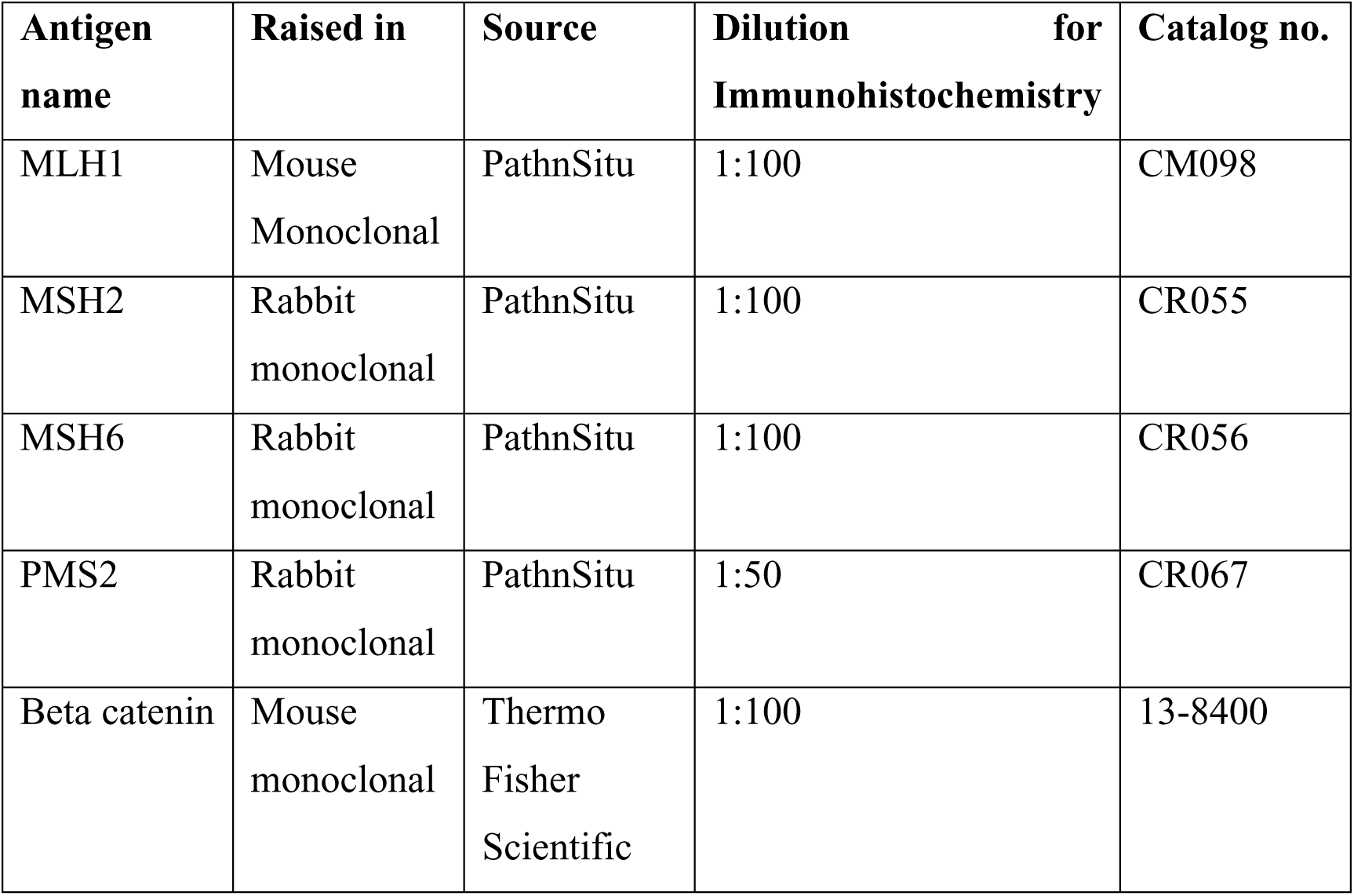

**List of primary antibodies used:**
| Antigen name | Raised in | Source | Dilution for Immunohistochemistry | Catalog no. |
| --- | --- | --- | --- | --- |
| MLH1 | Mouse Monoclonal | PathnSitu | 1:100 | CM098 |
| MSH2 | Rabbit monoclonal | PathnSitu | 1:100 | CR055 |
| MSH6 | Rabbit monoclonal | PathnSitu | 1:100 | CR056 |
| PMS2 | Rabbit monoclonal | PathnSitu | 1:50 | CR067 |
| Beta catenin | Mouse monoclonal | Thermo Fisher Scientific | 1:100 | 13-8400 |

### RNA isolation

RNA isolation was performed from fresh frozen colorectal (tumour and normal) tissues stored in RNALater at −80°C. Isolation was performed using the Qiagen AllPrep® DNA/RNA/miRNA Universal kit as per manufacturers’ protocol. Elution was done in Nuclease free water. For isolation of RNA from tissues in the validation cohorts, TRIzol reagent (Invitrogen) was used as per standard protocol.

### DNA isolation from tissue

DNA isolation from tissue was done by standard protocol. Chopped 50 mg tissue sample were incubated in digestion buffer (60mM Tris pH 8.0, 100mM EDTA, 0.5% SDS) and proteinase K (500ng/ml) at 56°C overnight. Equal volume of Phenol-chloroform solution was added and mixed well by inverting repeatedly. Tubes were centrifuged at 12,000 g for 15 minutes at room temperature for phase separation. Equal amount of chloroform was added and mixed well before centrifugation at 12000g for 15 minutes at room temperature for phase separation. 1/10th volume of 3M Sodium acetate pH 6.0 and 2.5 volumes of absolute ethanol were added to the aqueous phase and kept at −20°C overnight for nucleic acid precipitation. Centrifugation was done at 12500 g for 15 minutes at 4°C and washed with 70% ethanol at 12500g for 5 minutes at 4°C. The DNA pellet was air-dried and resuspended in 100µl of TE pH 8.0. RNase treatment (20µg/ml) was done for 30 minutes at room temperature to eliminate RNA. Phenol-chloroform phase separation and chloroform phase separation steps were repeated to remove the RNase and then DNA was purified from the aqueous phase by using Promega kit protocol (Promega Wizard SV Gel and PCR Clean-Up System, Part# 9FB072). Isolated DNA was quantitated by spectrometry and visualized by electrophoresis in 0.8% agarose gel using BioRad GEL DOC imaging system.

### PCR for APC gene

Normal and tumour DNA from each patient was subjected to PCR amplification. Primer sequences were designed to amplify the mutation cluster region (exon 15, codons 1260-1596) of the APC gene in overlapping PCR segments. Reaction conditions were as follows: 95°C, 5minutes; 95°C, 30 seconds; 60°C/ 57°C/60°C (Ta), 1 minute; 68°C, 1 minute; 68°C, 5 minutes; for 35 cycles. PCR amplification was done in a 25µl reaction with 1 unit of NEB Taq DNA polymerase (NEB Catalog# M0273S), 1x Standard Taq Buffer, 200µM dNTPs (NEB), 1µM forward and reverse primer each and 100 ng of template DNA. PCR product was visualized by electrophoresis in a 2% agarose gel in 1xTBE (0.13 M Tris (pH 7.6), 45 mM boric acid, 2.5 mM EDTA) buffer.

### Direct DNA sequencing

The PCR products were purified using the Promega Wizard SV Gel and PCR Clean-Up System according to the manufacturer’s instructions. Direct sequencing was performed by Eurofins Genomics India Pvt. Ltd.

### APC gene mutation analysis

The nucleotide and deduced amino acid sequences were compared with reference sequences of the APC gene available at the NCBI (National Center for Biotechnology Information) GenBank database using the BLASTx (Basic Local Alignment Search Tool) program.

### miRNA Seq Analysis

A total of 2-μg RNA sample isolated from 10 tissue types (5-tumour, 5-normal) for RNA sequencing. RNA samples were outsourced to the company, Bencos Research Solutions, for RNA sequencing and data analysis. RNA with RIN > 8.0 was proceeded for library preparation using the NEBNext® Small RNA Library Prep Set for Illumina® and sequenced at the National Genomics Core, CDFD, Hyderabad using a 50-bp single-end reads Illumina NextSeq 500 Sequencer (Illumina Inc., San Diego, CA, USA). The data was generated by using the paired end approach of Illumina technique. Total twenty paired end fastq files were used for the small RNA-seq analysis via a pipeline FastQC-FastpSortMeRNA-miRDeep2-edgeR. Fastq files were subjected to Fastqc (v.0.11.9) for the sequence quality check (FastQC; https://www.bioinformatics.babraham.ac.uk/projects/fastqc/) and found that all the quality features were passed, except some features which were flagged with warnings and failed. After sequencing, adapters and low-quality sequences were removed from obtained raw reads by the Fastp tool (v.0.23.2) (**12**) and clean data was counted using the FastQC program. All the reads of every sample were subjected to SortMeRNA (v.4.3.2) for removal of ribosomal RNA sequences (**13**). Four data bases, silva-euk-28s-id98, silva-euk-18s-id95, rfam-5.8s-database-id98 and rfam-5s-database-id98 were utilized in the rRNAs removal analysis (https://github.com/biocore/sortmerna/archive/2.1.tar.gz). Ten independent read mappings of small RNA-seq data were performed by miRDeep2 (v.2.0.1.2) using a genome reference sequence data base Homo_sapiens.GRCh38.dna.primary_assembly.fa (**14**) along with mature and hairpin miRNAs sequences specific to human (hsa as a species) retrieved from the miRBase (v22.1) database (https://www.mirbase.org/ftp.shtml). The reference sequence fasta file was downloaded from the Ensembl genome database (http://ftp.ensembl.org/pub/release-105/fasta/homo_sapiens/dna/).

### Identification of known and novel miRNAs

Processed reads were used to generate collapsed reads using mapper.pl module of the miRDeep2 package with minimum length 18 parameter. For predicting miRNAs, known and novel, the collapsed reads were passed to miRDeep2.pl module of the package. In this analysis, the reference genome sequences and the miRBase mature and hairpin sequences specific to human (hsa as a species) were utilized. Count matrix was generated using the final results after removing the duplicates based on the same genomic coordinates.

### Differential miRNA expression analysis

Read count was performed for all the samples using the miRDeep2. For differential miRNA expression analysis, edgeR (v.3.36.0) R-package was utilized (**15**). Normalization factor was calculated using raw read counts, and after that the count data was normalized by the Count Per Million (CPM) method. The exact Test method available in the egdeR was implemented for the differential expression analysis (DEA) of the single sample comparisons. In the single sample comparisons, 0.2 divergence value was considered for the DEA. However, lmFit module of the edgeR, a linear model using weighted least squares for each gene, was utilized to fit the linear model in to the data for multiple samples comparison after applying the voom transformation and calculation of variance weights. Further, empirical Bayes (eBayes) module of the edgeR was employed for smoothing the standard errors and calling the differentially expressed transcriptsmiRNAs. Differentially expressed miRNAs were obtained by filtering the results of DEA using adjusted p-value or FDR ≤ 0.05 and Log_2_ Fold Change value at ≥2 (upregulation) or ≤ −2 (downregulation).

### Volcano Plots

Global gene expression values obtained from a pair-wise comparison analysis was also plotted in the form of volcano plot using the R-package. In the volcano plots, miRNAs rows were ordered from the final result of edgeR analysis according to the adjusted p-value or FDR in decreasing order. The volcano plot was generated for all the five single sample comparisons.

### Heatmap

The pheatmap package in R was implemented to plot the heatmaps, and the rlog-normalised read count matrix for all the samples was used as the input data (https://cran.r-project.org/web/packages/pheatmap/index.html) derived from the DESeq2 R-package (**16**). The function rlog returns a Summarized Experiment object which contains the rlog-transformed values in its assay slot. Corresponding Z-scores were computed from the rlog-normalized read count matrix, and pheatmap drew the heatmap accordingly. The top 20 most variable miRNAs were extracted from the matrix to be plotted onto the heatmap. The miRNAs that showed expression values higher than the mean expression across samples were assigned a positive Z-score denoted by green. Whereas the opposite, that is, negative Z-score is denoted by a red colour on the heatmap.

### Target identification and selection

Experimentally validated targets for the selected miRNAs were identified using miRNet (**17**). Additionally, genes differentially expressed (DE) in TCGA colorectal cancer (COAD) samples with |Log2FC| cutoff of 1.00 and q-value cutoff of 0.01 were derived using GEPIA (**18**). Target genes were compared with this DE-gene list to identify DE-targets altered in colorectal tumour tissue in a direction reciprocal to that of miRNAs.

### Gene Ontology and Pathway Enrichment Analysis

For each comparison group, gene ontology (GO) and pathway enrichment analysis were performed separately for both the upregulated and downregulated set of differentially expressed miRNAs’ target genes against human as a selected species. A tool ShinyGO (v.0.75) (**19**) was used for retrieving functional annotations based on the Biological Process (BP), Molecular Function (MF) and Cellular Component (CC). FDR was calculated based on nominal p-value from the hypergeometric test.

### miRNA validation by qRT-PCR

Real time analyses by two-step RT-qPCR was performed for quantification of miRNA levels. The stem loop qRT-PCR method was used for miRNA screening and quantification (**20**, **21**). Reverse transcription (RT) was performed with Verso cDNA Synthesis Kit (Thermo Scientific #AB-1453/A) as per the manufacturer’s instructions using 100ng of total cellular RNA. The 10µl of RT reaction mixture contained 1 µl of RT primer (1 µM), 500µM each of dNTP, 2 µl of 5x cDNA synthesis buffer, 0.5 µl of RT enhancer and 0.5 µl of Verso Enzyme Mix. All miRNA RT-qPCRs were performed on the Biorad CFX96 Real Time System. One tenth of the reverse transcription mix was subjected to PCR amplification with Bio-Rad SsoAdvanced Universal SYBR® Green Supermix (Catalog# 1725270). The RT reaction condition was: 25°C, 10 min; 42°C, 60 min; 95°C, 5 min; 4°C, ∝. The PCR condition was: 95°C, 3 min; 95°C, 30 sec; 60°C, 1 min; for 40 cycles. All Samples were analyzed in triplicates. The concentrations of intra cellular miRNAs were calculated based on their normalized Ct values. Normalization was done by U6 snRNA. The ΔΔCt method for relative quantitation (RQ) of gene expression was used and relative quantification was done using the equation 2-ΔΔCt (as per ‘Guide to Performing Relative Quantitation of Gene Expression Using Real-Time Quantitative PCR’ obtained from the Applied Biosystems website (http://www3.appliedbiosystems.com/cms/groups/mcb_support/documents/generaldocuments/c ms_042380.pdf).

### Statistical Analysis

All graphs were plotted and analysed using GraphPad Prism 5.00 (GraphPad, San Diego, CA, USA). For statistical analysis, non-parametric two-tailed, paired or unpaired Student’s t-test was performed. Error bars indicate mean with standard deviation.

Information on PCR primers, miRNA Reverse Transcription (RT) Stem Loop Primers (SLP), miRNA Real time PCR forward and reverse primers and Antibodies are provided below:

### List of PCR primers

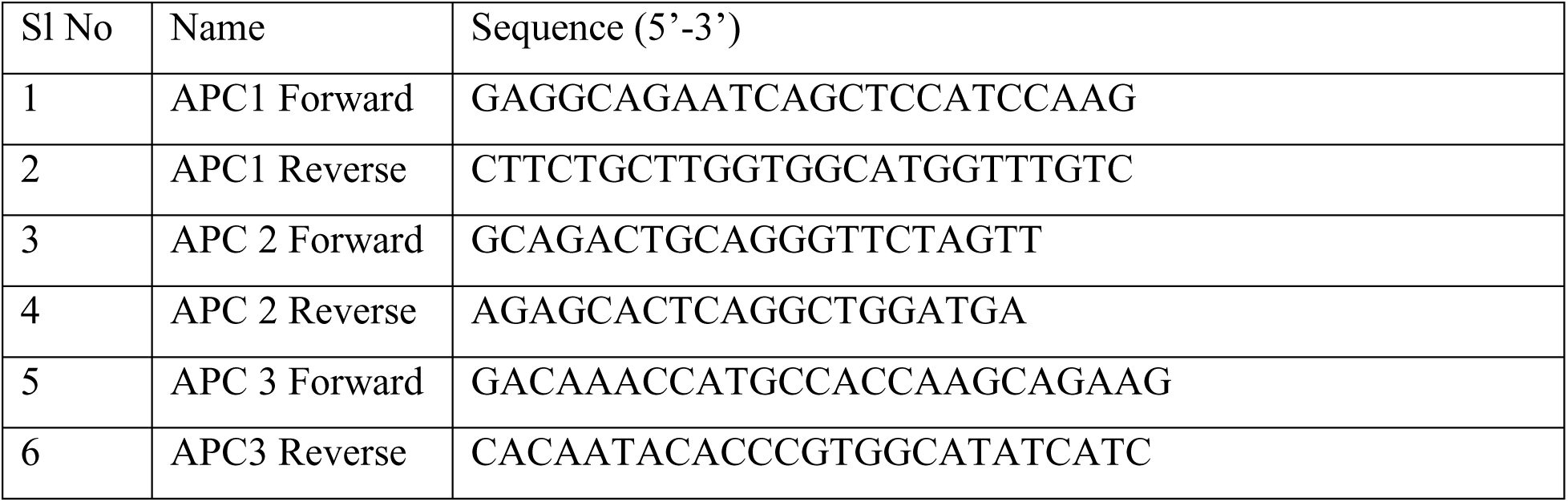

Primer sequences for Reverse Transcription (RT) Stem Loop Primers (SLP) used in study.

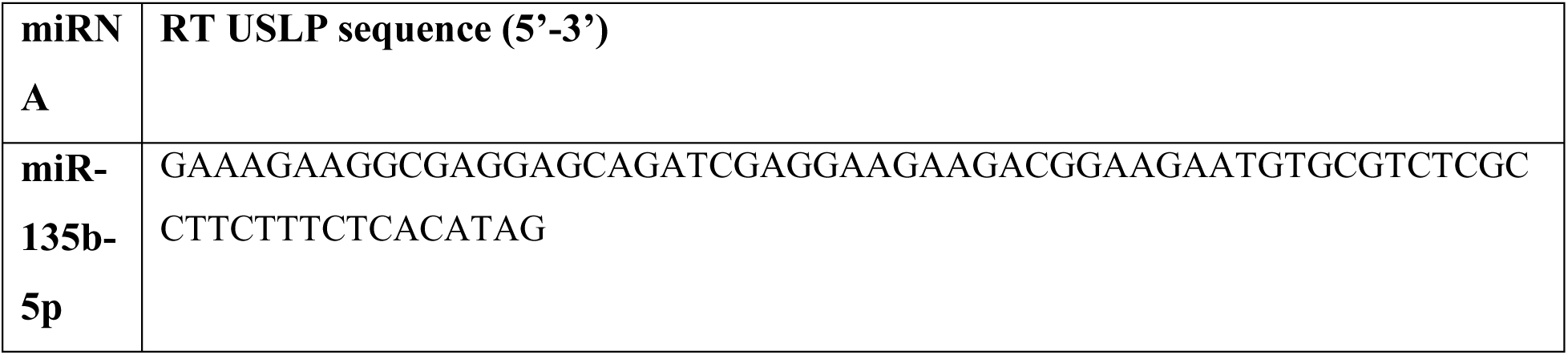
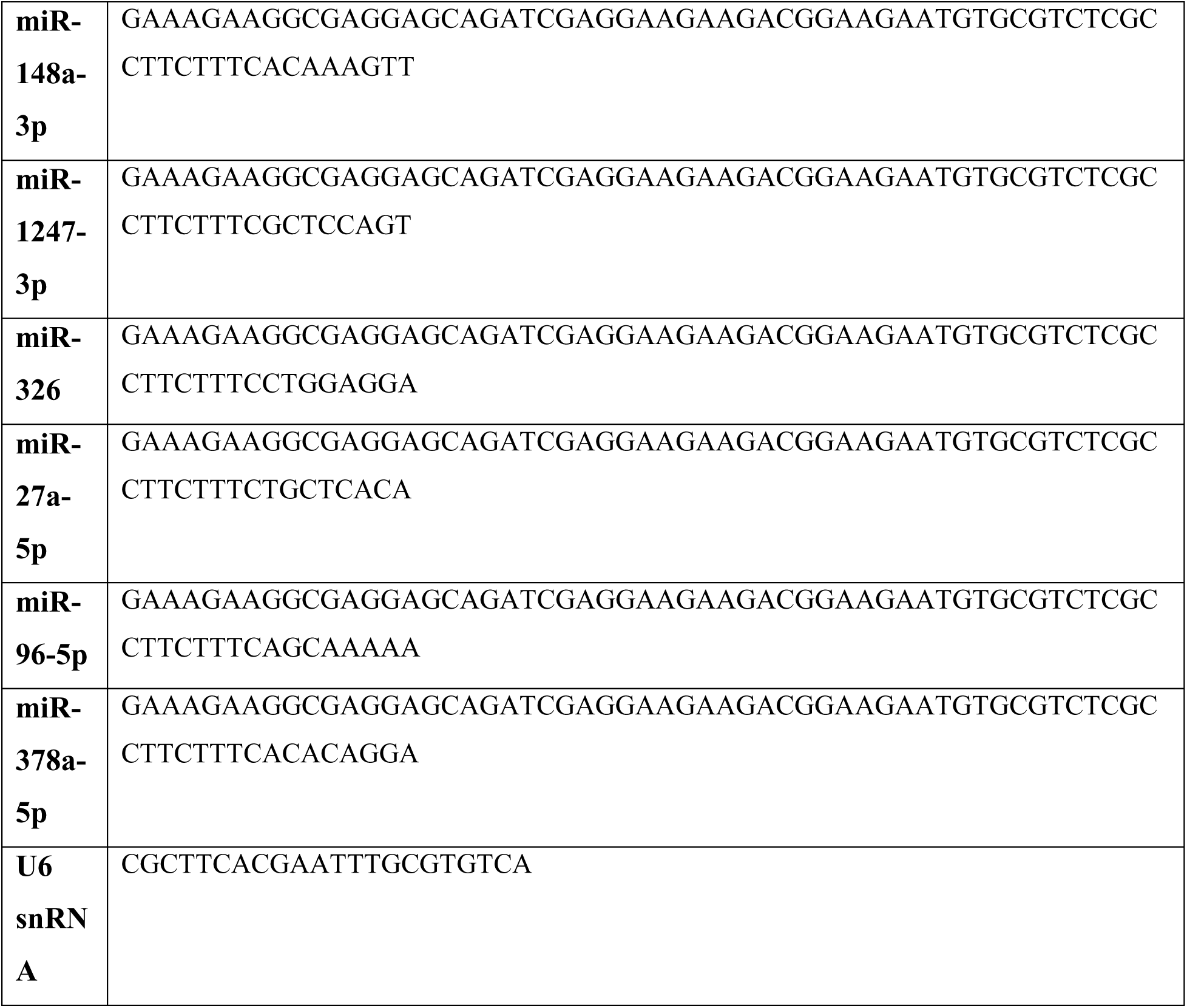

Primer sequences for Real time PCR Forward and reverse primers. The reverse primer sequence is complementary to a portion of the RT USLP and is hence common for all miRNAs.

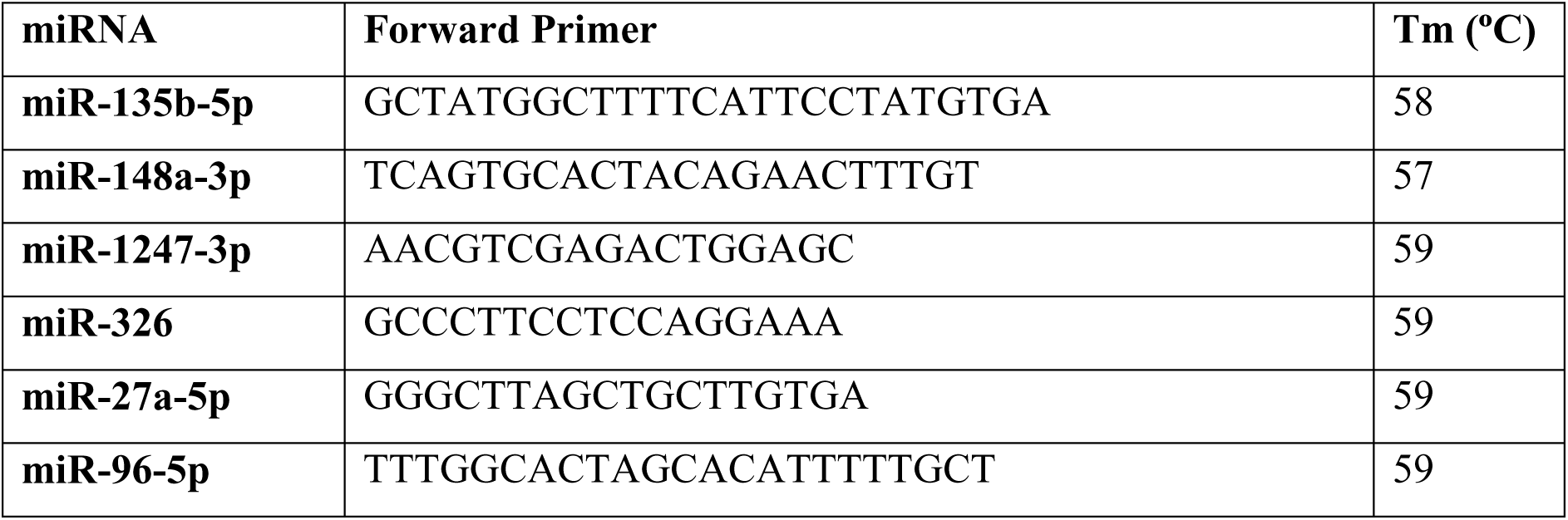
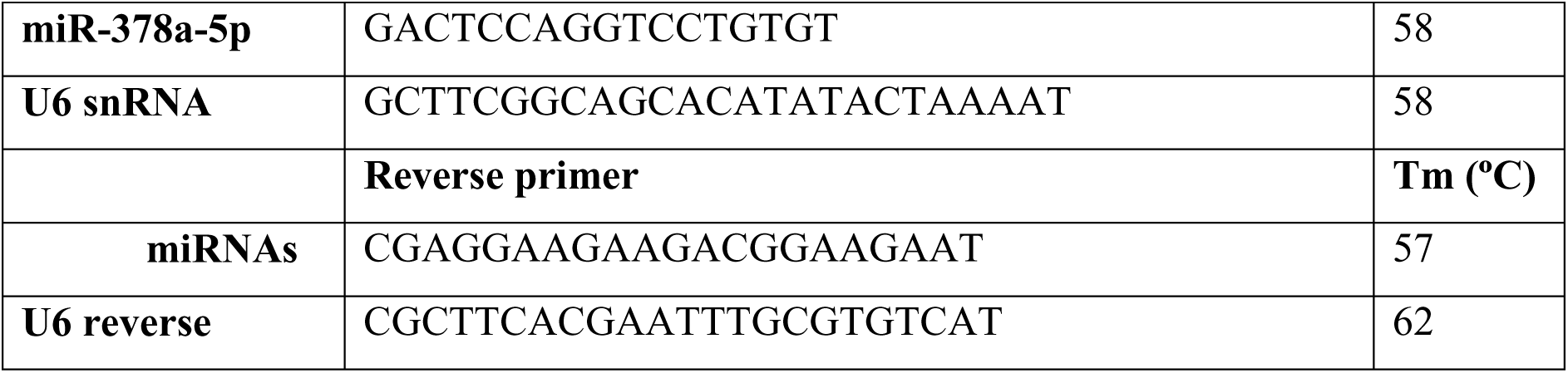

## Results

### Patient recruitment and sample collection

34 histologically confirmed colorectal adenocarcinomas and adjacent colorectal mucosas were collected. Upon histopathological analysis by a trained pathologist, malignant lesions were found to be of the following TNM stages: T1 (11.76%), T2 (29.41%), T3 (41.2%) and T4 (17.65%). 11.76% of them represented well differentiated adenocarcinoma (Grade: G1), 70.6% were moderately differentiated (Grade: G2), and 14.71% were poorly differentiated (Grade: G3). Screening was done to eliminate tumours with Micro Satellite Instability (MSI), nuclear localization of β-catenin and *APC* mutations (Mutation Cluster Region-between codons 1260 and 1596 of exon 15 of the APC gene). Representative images of immunohistochemical detection of MSI and nuclear β-catenin have been shown in Supplementary Figures S1 and S2 respectively.

### NGS small-RNA sequencing and analysis

Out of the patients who were Micro Satellite Stable, without nuclear β-catenin and lacking somatic *APC* mutations, tumours and paired normal tissues of 5 (Patient no. 1, 2, 3, 4 and 5) were sent for NGS based miRNA sequencing. The 5 patients which were chosen for miRNA seq consisted of 3 young patients (mean age: 43 years) and 2 old patients (mean age: 63 years). Pathological information of the clinical samples sent for miRNA-seq have been summarized in Table 1.

**Table 1:** Clinicopathological features of clinical samples (Tumour and Paired normal from each patient) sent for NGS based miRNA Sequencing (Discovery cohort).

| Young/<br>Old | Patient<br>number | Age | Gender | Stage | Grade | Histopathological<br>type | Location |
| --- | --- | --- | --- | --- | --- | --- | --- |
| Young | 1 | 39 | Female | T<sub>3</sub>N<sub>2b</sub>M<sub>x</sub> | G2 | Adenocarcinoma | Junction of ascending<br>and transverse colon |
| Young | 2 | 44 | Female | T<sub>4a</sub>N<sub>0</sub> | G2 | Adenocarcinoma | Rectosigmoid colon |
| Young | 3 | 47 | Male | T<sub>2</sub>N<sub>0</sub> | G2 | Adenocarcinoma | Ascending colon and<br>cecum |
| Old | 4 | 66 | Male | T<sub>3</sub>N<sub>2b</sub> | G2 | Adenocarcinoma | Rectum |
| Old | 5 | 60 | Female | T<sub>1</sub>N<sub>0</sub> | G1 | Adenocarcinoma | Rectum |

Sequencing data of the ten human samples (normal and tumour tissue of 5 CRC patients) were obtained for the small RNA (miRNA) transcriptome analysis. Total twenty paired end fastq files were used for the small RNA-seq analysis via a pipeline FastQC-Fastp-SortMeRNA-miRDeep2-edgeR. The schematic workflow for the overall analysis has been depicted in Figure 1.

**Figure 1.**
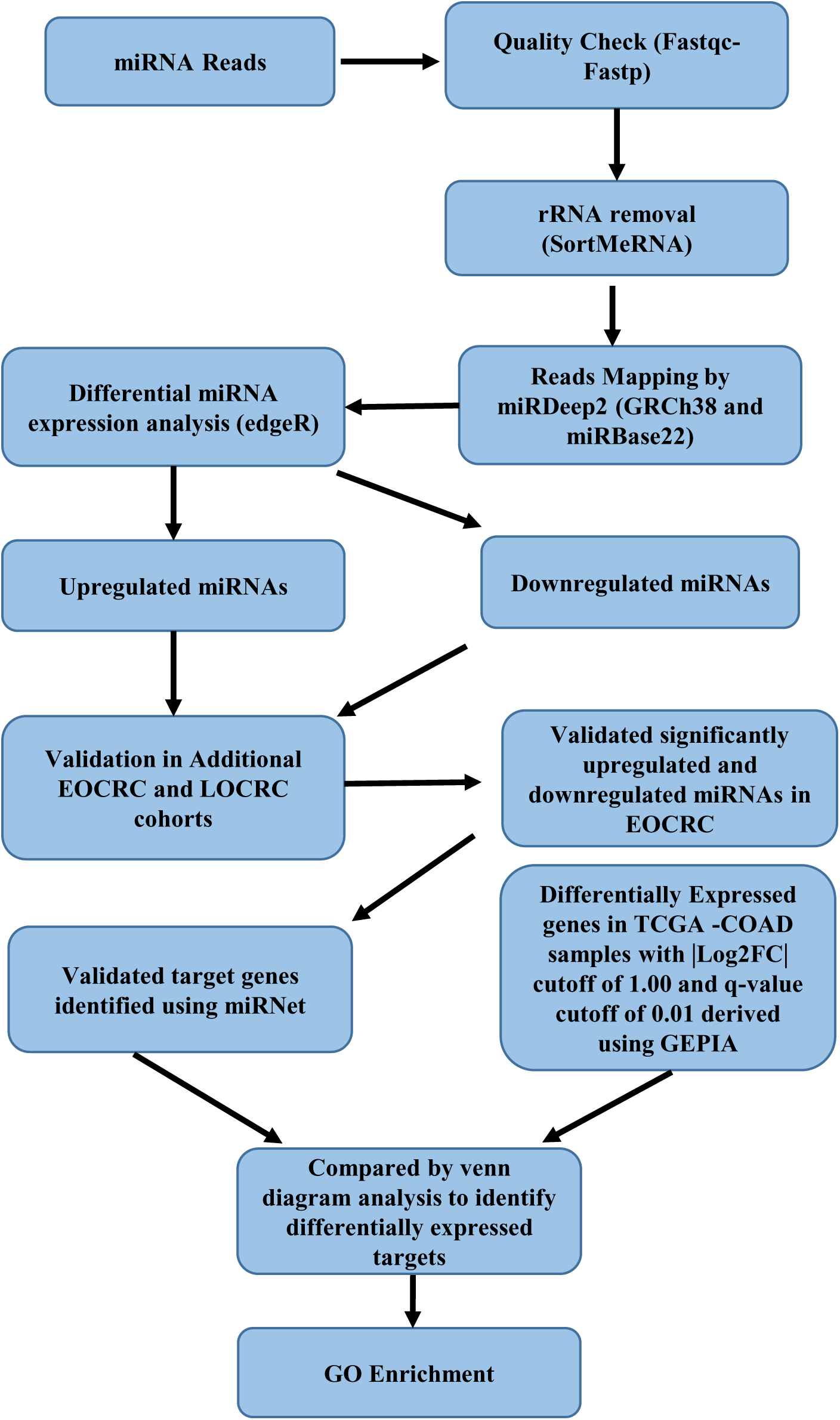
Schematic work-flow of overall pipeline used for the small RNA (miRNA) transcriptome analysis. Schematic work-flow of overall pipeline used for the small RNA (miRNA) transcriptome analysis of sequencing data generated from 5 normal colonic mucosas and 5 colorectal tumours. Total twenty paired end fastq files were used for the small RNA-seq analysis via a pipeline FastQC-FastpSortMeRNA-miRDeep2-edgeR. Top differentially expressed miRNAs were validated in additional Early-Onset Colorectal Carcinoma (EOCRC) and Late-Onset Colorectal Carcinoma (LOCRC) cohorts. Validated targets of significantly differentially expressed miRNAs were compared with differentially expressed genes in TCGA-COAD datasets to identify potential targets deregulated in EOCRC.

Sequencing the miRNA libraries for CRC tissues and paired normal tissues resulted in a total of 91,658,630 and 94,627,488 raw reads, respectively. The removal of adaptor sequences, junk reads, reads other than 15 to 30 bp, rRNA, snRNA, snoRNA, and tRNA produced 83,719,335 and 90,030,493 clean reads respectively. The summary of read alignment has been indicated in Supplementary Table 1. Atleast 85% of the raw reads accounted for the clean reads, which suggested that a useful group of miRNAs was obtained with a reasonable sequencing depth. Overall mapping rate was over 28% among samples. Read depth coverage and sequence length distribution plots have been depicted in Supplementary Figure S3.

### Differential miRNA expression analysis

For single sample comparisons, normal colonic mucosa of each patient was taken as the normal control for differential expression analysis. The dataset for the normal control of each patient was compared with the tumour data set of that patient. For multiple sample comparisons, the patient datasets were divided into two groups-young and old based on their age. Patients 1, 2 and 3 were considered as young as their age was < 50 years and patients 4 and 5 were considered as old as their age was > 55 years. In the Young Normal vs Tumour comparison, the normal dataset was compared with the tumour dataset of young patients (normal as control group and tumour as test group). Similarly, in the Old Normal vs Tumour comparison, the normal dataset for old patients was compared with the tumour dataset for the same (normal as control group and tumour as test group). Differentially expressed miRNAs (DEMs) were obtained by filtering the results of DEA using p-value <0.05 and Log2 Fold Change value at >2 (upregulation) or < −2 (downregulation). The volcano plots for each sample comparison of each patient has been depicted in <u>Figure 2 A-E</u>. These were generated having cut-off criteria log2fold change ≥2 (p<0.05) for up regulated and ≤−2 (p<0.05) for down regulated genes.

**Figure 2.**
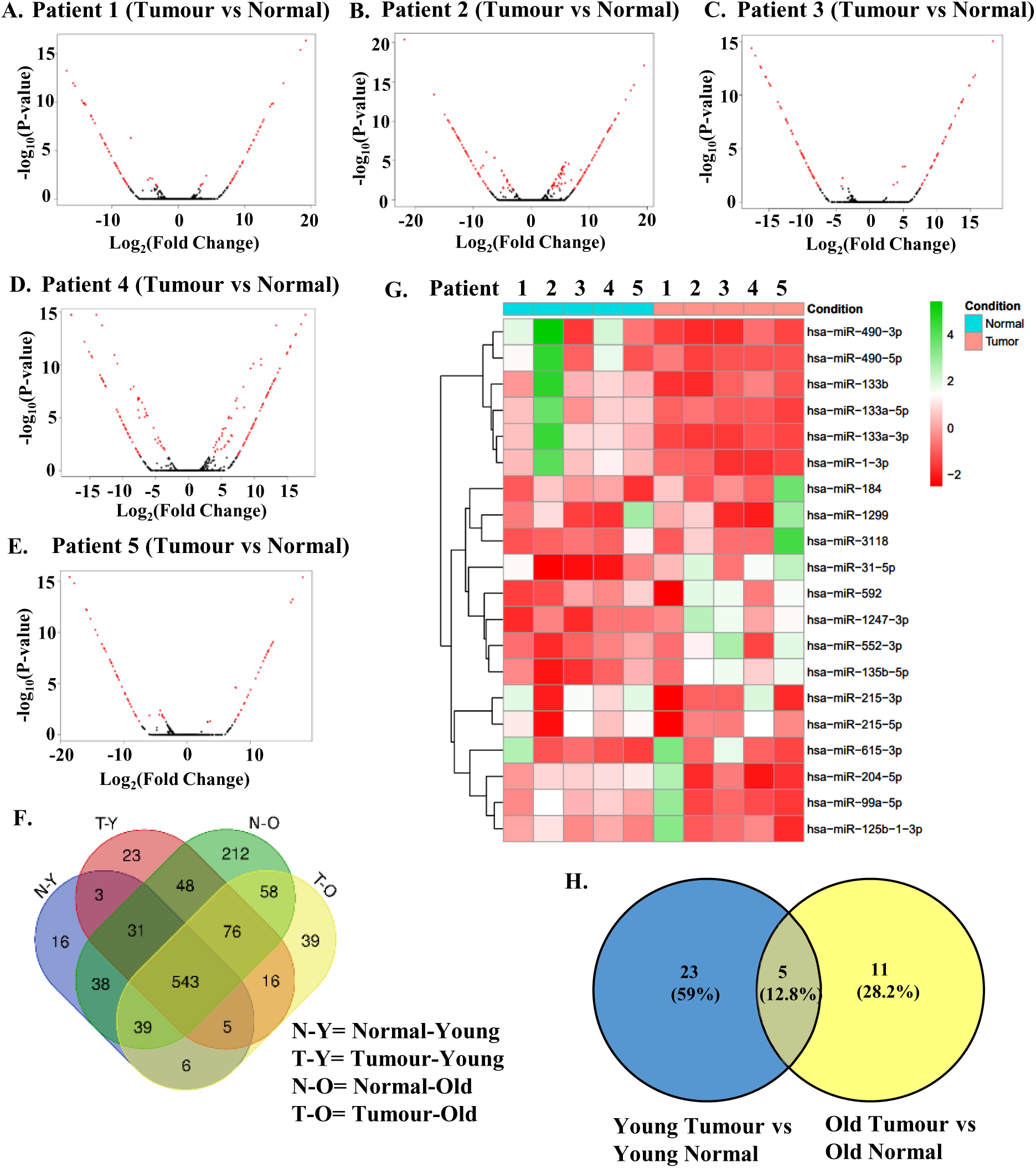
Identification of Differentially Expressed miRNAs (DEMs) in tumor samples of the discovery cohort. **(A-E)**: Significant differentially expressed miRNAs were identified by filtering the results with a cut off parameter adjusted p-value or FDR <0.05 Global miRNA expression values obtained from a pair-wise comparison (Tumour miRNA compared with corresponding normal) analysis for each patient in the discovery cohort was plotted in the form of volcano plots. Differentially expressed miRNAs at the level of statistical significance adjusted p-value (FDR) <0.05 are shown in red colored dots, while non-significant miRNAs are in black colored dots. X-axis shows negative logarithm of FDR values, and y-axis represents log fold change value of each transcript. Patients 1, 2 and 3 represent young patients <50 years old, Patients 4 and 5 represent old patients >55 years old **(F)** All identified miRNAs of Normal_old, Tumor_old, Normal_young, Tumor_young were used for the generation of venn diagram. The venn diagram was constructed via Venn (https://bioinformatics.psb.ugent.be/webtools/Venn/). **(G)** Heatmap of top 20 miRNAs sorted by variance in decreasing order, found across 10 samples. The heatmap was generated by using rlog-normalized count data of each sample. Z-scores for each miRNA per sample are indicated in red and green colour scale shown on the right, for expression levels below and above mean expressions across sample for that given miRNA, respectively. The Condition legend denotes the sample types, where cyan blue is assigned to Normal samples and Pink is assigned to Tumor samples **(H)** Venn diagram showing known DEMs with cut-off p-value <0.05 and Log2 Fold Change value at >2 or < −2 identified in the multiple sample comparisons of Young Tumour vs Normal and Old Tumour vs Normal (normal as control group and tumour as test group). Constructed using https://bioinfogp.cnb.csic.es/tools/venny/.

All Identified miRNAs (both known and novel) of Normal_old (N-O), Tumor_old (T-O), Normal_young (N-Y), Tumor_young (T-Y) were used for the generation of venn diagram (Figure 2F). Known DEMs with cut-off p-value <0.05 and Log2 Fold Change value at >2 or < −2 identified in the multiple sample comparisons of Young Tumour vs Normal and Old Tumour vs Normal (normal as control group and tumour as test group) were analysed by venn diagram (https://bioinfogp.cnb.csic.es/tools/venny/) (Figure 2H). Heatmap of the top 20 miRNAs (Figure 2G) and all miRNAs (Supplementary Figure S4) identified by RNA-Sequencing of our 10 samples and sorted by variance in decreasing order was generated. 23 DEMs were identified specific to the Young Tumour vs Normal group whereas 11 were identified unique to the Old Tumour vs Normal group. 5 DEMs were common to both groups indicating overall differential expression in CRC. The 5 common elements differentially expressed in both subsets were-hsa-miR-129-5p, hsa-miR-9-5p, hsa-miR-1-3p, hsa-miR-145-5p and hsa-miR-133a-3p. The full list of DEMs identified in both young and old patients have been described in Supplementary Table 2. The list of identified DEMs specific to young tumours has been given in Table 2.

**Table 2:** List of significant differentially expressed miRNAs identified in our bioinformatics analysis of miRNA-sequencing in young patient tumours compared with corresponding normal (DEM cut-off: p-value <0.05, Log Fold Change: >2, <-2)

| miRNA | logFC | AveExpr | P-Value | Upregulated/<br>Down regulated |
| --- | --- | --- | --- | --- |
| hsa-miR-1247-3p | 3.70881538 | 2.13482567 | 0.00174089 | Up |
| hsa-miR-27a-5p | 2.61304708 | 7.28367881 | 0.00901648 | Up |
| hsa-miR-96-5p | 2.80991462 | 6.01862554 | 0.02211184 | Up |
| hsa-miR-148a-3p | 2.49604406 | 1.99822506 | 0.02365696 | Up |
| hsa-miR-135b-5p | 3.41150654 | 6.19388937 | 0.02799316 | Up |
| hsa-miR-133b | -2.01434615 | 1.8466954 | 0.00292072 | Down |
| hsa-miR-133a-5p | -2.13259517 | 0.22441615 | 0.00324305 | Down |
| hsa-miR-378e | -2.61154969 | 1.13114207 | 0.00366325 | Down |
| hsa-miR-139-3p | -2.1809868 | 1.433196 | 0.00568353 | Down |
| hsa-miR-378a-5p | -2.41712807 | 3.98406158 | 0.00620528 | Down |
| hsa-miR-887-3p | -2.10734608 | 0.13910073 | 0.00627189 | Down |
| hsa-miR-30a-5p | -2.0469516 | 9.41801359 | 0.00769662 | Down |
| hsa-miR-9-3p | -2.15316894 | 1.64216281 | 0.00770244 | Down |
| hsa-miR-490-3p | -2.11550509 | 1.30918108 | 0.01130573 | Down |
| hsa-miR-143-3p | -2.29934116 | 15.5726929 | 0.01331962 | Down |
| hsa-miR-378c | -2.49476989 | 7.31380586 | 0.01533526 | Down |
| hsa-miR-490-5p | -2.14003278 | 0.29238366 | 0.01709929 | Down |
| hsa-miR-326 | -2.63728355 | 1.68062835 | 0.01712207 | Down |
| hsa-miR-504-5p | -2.28494574 | 3.64916906 | 0.01904966 | Down |
| hsa-miR-378d | -2.48909916 | 6.12055306 | 0.01914212 | Down |
| hsa-miR-143-5p | -2.0070137 | 7.68776446 | 0.02078392 | Down |
| hsa-miR-139-5p | -2.27780576 | 6.20200982 | 0.0240824 | Down |
| hsa-miR-363-3p | -2.40777269 | 6.41529968 | 0.04421748 | Down |

### Identification of DEMS to be validated in additional cohorts based on TCGA data analysis

Out of the 23 DEMs specific to the EOCRC subset, we chose the top 10 DEMs (on the basis of LogFC) for further validation using TCGA datasets. The top 10 miRNAs differentially expressed in young patient tumours of the discovery cohort in our small-RNA sequencing analysis were hsa-miR-1247-3p, hsa-miR-27a-5p, hsa-miR-96-5p, hsa-miR-326, hsa-miR-378c, has-miR-378d, hsa-miR-378a-5p, hsa-miR-378e, hsa-miR148a-3p and hsa-miR-135b-5p (Table 2). These miRNAs were tested on the TCGA dataset of colorectal adenocarcinomas (COAD) using matched TCGA normal dataset (<u>CancerMIRNome (jialab-ucr.org)</u>) (**22**). We observed significant and constant differential expression of hsa-miR-1247-3p, hsa-miR-27a-5p, hsa-miR-96-5p, hsa-miR-326, hsa-miR-378a-5p, hsa-miR148a-3p and hsa-miR-135b-5p among large data sets (Figure 3 A-G). Upregulation or downregulation observed was similar to that obtained by our NGS analysis. Hsa-miR-378e, hsa-miR-378c and hs-miR-378d did not show significant differential expression in the TCGA dataset and were therefore not selected for further validation in additional patient (young/ aged) cohorts (Figure 3 H-J).

**FIGURE 3.**
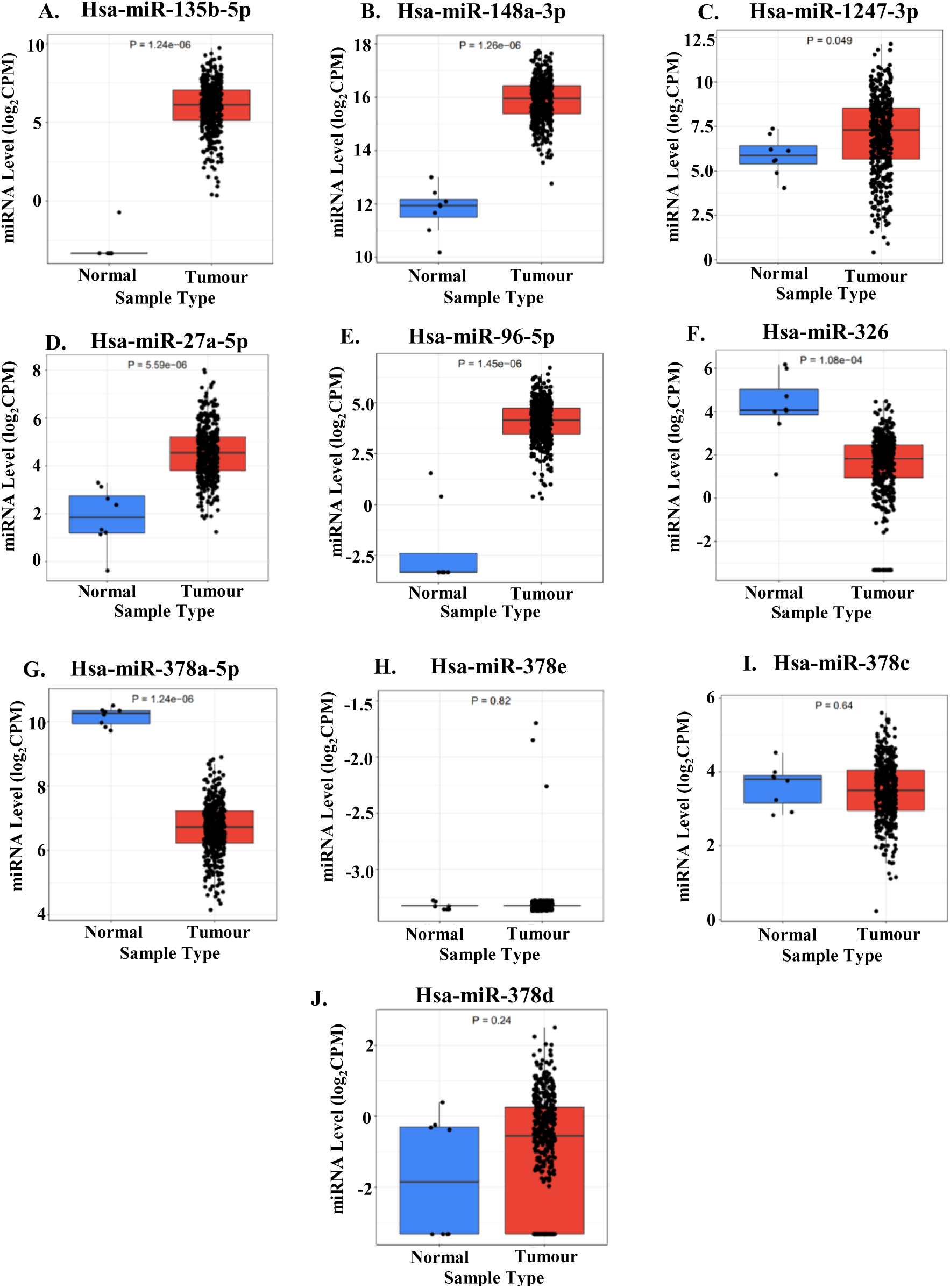
TCGA based validation of top 10 miRNAs differentially expressed in sample comparison of young tumours with young normal tissues on Cancer MIRNome (CancerMIRNome (jialab-ucr.org)). Gene expression analysis of **(A)** hsa-miR-135b-5p, **(B)** hsa-miR-148a-3p, **(C)** hsa-miR-1247-3p, **(D)** hsa-miR-27a-5p, **(E)** hsa-miR-96-5p, **(F)** hsa-miR-326, **(G)** hsa-miR-378a-5p, **(H)** hsa-miR-378e, **(I)** hsa-miR-378c and **(J)** hsa-miR-378d on the TCGA dataset of colorectal adenocarcinomas (COAD).

### Validation of selected DEMs in EOCRC

We validated the selected miRNAs significantly differentially expressed in the TCGA-COAD dataset (hsa-miR-1247-3p, hsa-miR-27a-5p, hsa-miR-96-5p, hsa-miR-326, hsa-miR-378a-5p, hsa-miR148a-3p and hsa-miR-135b-5p) by qRT-PCR in an additional EOCRC validation cohort of 10 young patients (<50 years old). Adjacent normal colonic mucosa of each tumour was used as the corresponding control. Table 3 details the clinicopathological details of patients, both EOCRC and LOCRC (Tumour tissue and Normal mucosa), included in the validation cohort. miRNA levels were detected by qRT-PCR using stem loop primers for cDNA synthesis and PCR amplification was done using specific forward and reverse primers (**20**, **21**).

**Table 3:** Clinicopathological features of clinical samples (Tumour and Paired normal from each patient) used for validation of NGS results.

| **Young/**<br>**Aged** | **Patient**<br>**number** | **Age** | **Gender** | **Stage** | **Grade** | **Histopathological**<br>**type** | **Location** |
| --- | --- | --- | --- | --- | --- | --- | --- |
| Young | Y1 | 27 | Male | ycT<sub>4a</sub>N<sub>2</sub>M<sub>0</sub> | G2 | Adenocarcinoma | Ascending colon |
| Young | Y2 | 25 | Male | cT<sub>4</sub>N<sub>2a</sub> | G3 | Adenocarcinoma | Lower Rectum |
| Young | Y3 | 42 | Male | cT<sub>3</sub>N<sub>0</sub>M<sub>0</sub> | G2 | Adenocarcinoma | Transverse colon |
| Young | Y4 | 39 | Male | cT<sub>3</sub>N<sub>2</sub> | G1 | Adenocarcinoma | Transverse colon |
| Young | Y5 | 11 | Male | cT<sub>3</sub>N<sub>1</sub>M<sub>0</sub> | G3 | Adenocarcinoma | Transverse colon |
| Young | Y6 | 18 | Female | T<sub>4b</sub>N<sub>2</sub>M<sub>1b</sub> | G3 | Adenocarcinoma | Upper rectum |
| Young | Y7 | 32 | Male | cT<sub>3</sub>N<sub>0</sub>M<sub>0</sub> | G2 | Adenocarcinoma | Sigmoid colon |
| Young | Y8 | 45 | Female | cT<sub>4</sub>N<sub>0</sub>M<sub>1c</sub> | G1 | Adenocarcinoma | Upper rectum |
| Young | Y9 | 18 | Female | cT<sub>3</sub>N<sub>2</sub>M<sub>0</sub> | G2 | Adenocarcinoma | Lower rectum |
| Young | Y10 | 43 | Male | cT<sub>4a</sub>N | G2 | Adenocarcinoma | Distal rectum |
| Aged | O1 | 58 | Male | cT<sub>3</sub>N<sub>2</sub>M<sub>0</sub> | G2 | Adenocarcinoma | Ascending colon |
| Aged | O2 | 70 | Male | T<sub>4b</sub>N<sub>2</sub>M<sub>1b</sub> | G3 | Adenocarcinoma | Lower rectum |
| Aged | O3 | 61 | Male | cT<sub>3</sub>N<sub>2</sub>M<sub>0</sub> | G2 | Adenocarcinoma | Ascending colon |
| Aged | O4 | 75 | Male | T<sub>3</sub>N<sub>2</sub>M<sub>0</sub> | G3 | Adenocarcinoma | Hepatic Flexure |
| Aged | O5 | 59 | Male | T<sub>2</sub>N<sub>0</sub>M<sub>0</sub> | G2 | Adenocarcinoma | Lower rectum |
| Aged | O6 | 63 | Male | T<sub>4</sub>N<sub>2b</sub>M<sub>1a</sub> | G2 | Adenocarcinoma | Rectosigmoid, |
| Aged | O7 | 57 | Female | cT<sub>3</sub>N<sub>1</sub>M<sub>0</sub> | G2 | Adenocarcinoma | Hepatic Flexure |

Similar expression pattern was observed for hsa-miR-1247-3p, hsa-miR-148a-3p, hsa-miR-135b-5p, hsa-miR-27a-5p, hsa-miR-96-5p and hsa-miR-326 in all 10 young patients supporting our NGS data analysis results (Figure 4 A-F). These miRNAs showed significant differential expression as compared to normal tissue in the EOCRC validation cohort. However, significant downregulation of hsa-miR-378a-5p (Figure 4 G) was not observed in the validation cohort.

**FIGURE 4.**
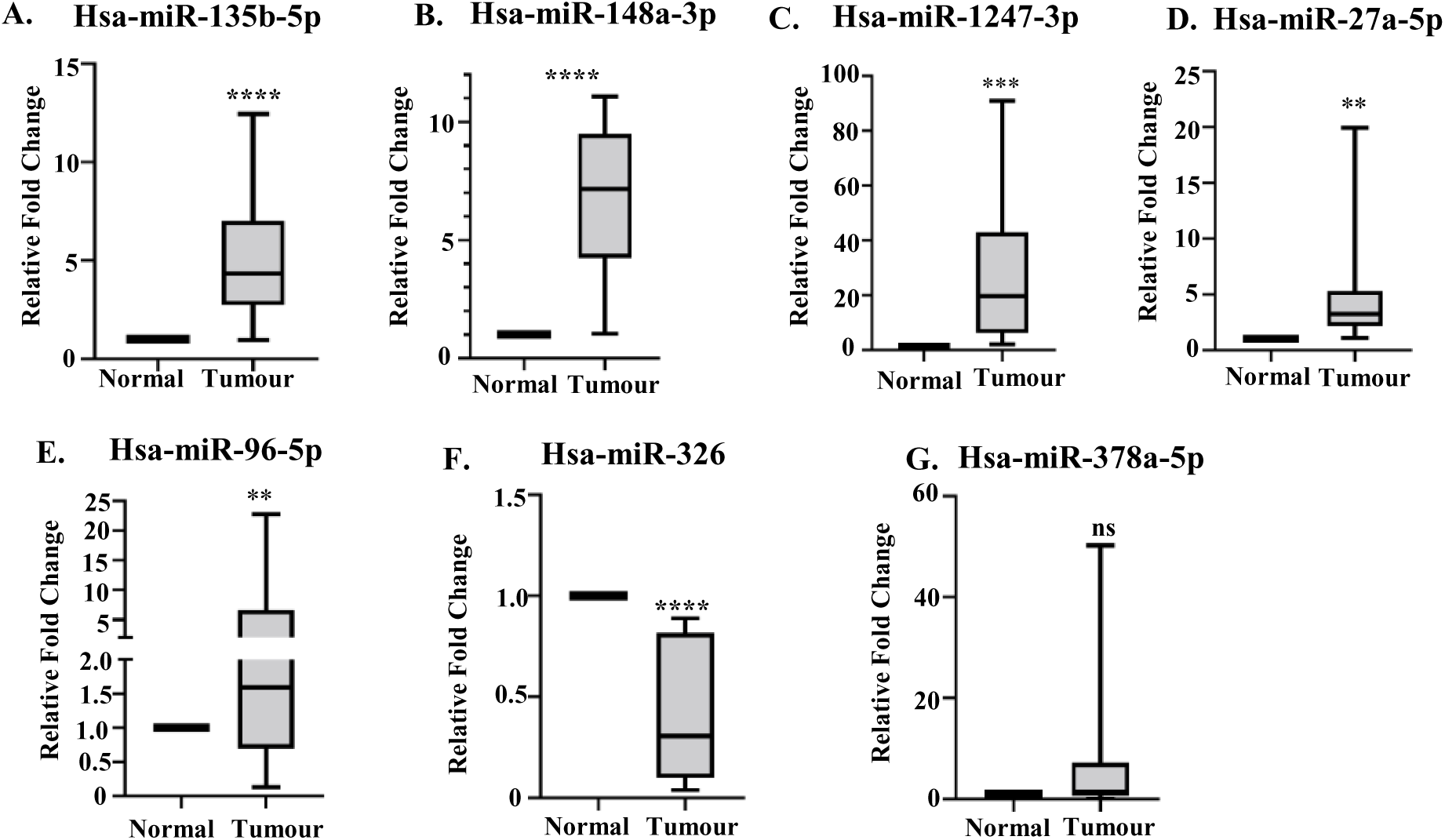
qRT-PCR based differential analysis of TCGA validated DEMs of young tumours vs young normal tissues in an additional EOCRC validation cohort. Relative expression analysis of **(A)** hsa-miR-135b-5p, **(B)** hsa-miR-148a-3p, **(C)** hsa-miR-1247-3p, **(D)** hsa-miR-27a-5p, **(E)** hsa-miR-96-5p, **(F)** hsa-miR-326 and **(G)** hsa-miR-378a-5p in tumours and normal tissues of 10 young patients (<50 years old) of our EOCRC validation cohort. RT-qPCR detection of miRNAs was done from 200 ng of isolated cellular RNA. N=3 replicates for each patient tissue (tumour and normal). Normalization was done by U6 snRNA. Data represents Mean ± SD. Statistical significance was calculated by two-tailed Students’ paired *t*-test. ns: non-significant, **P < 0.01, ***P < 0.001. ****P < 0.0001

To verify if differential expression of the above mentioned miRNAs was restricted to the young population, we cross-checked their expression in an additional validation cohort of 7 aged patients (LOCRC, ≥55 years old). Clinicopathological details of old patients included in the aged validation cohort have been given in Table 3. Figure 4 demonstrates the distribution of the selected DEMs in the young patient validation cohort and figure 5 shows their expression in the aged patient validation cohort. Hsa-miR-135b-5p, hsa-miR-148a-3p, hsa-miR-27a-5p and hsa-miR-96-5p were significantly upregulated compared to normal, both in the young and aged patient validation cohort (Figure 4 A, B, D, E and 5 A, B, D, E). Thus these miRNAs do not show age-specific expression. Upregulation of hsa-miR-1247-3p and downregulation of hsa-miR-326 was found to be significant only in the young validation cohort (Figure 4C, 4F and 5C, 5F). These findings suggest that hsa-miR-1247-3p and hsa-miR-326 are differentially expressed in EOCRC. Understanding their regulatory pathways may indicate potential therapeutic avenues to be targeted in EOCRC.

**FIGURE 5.**
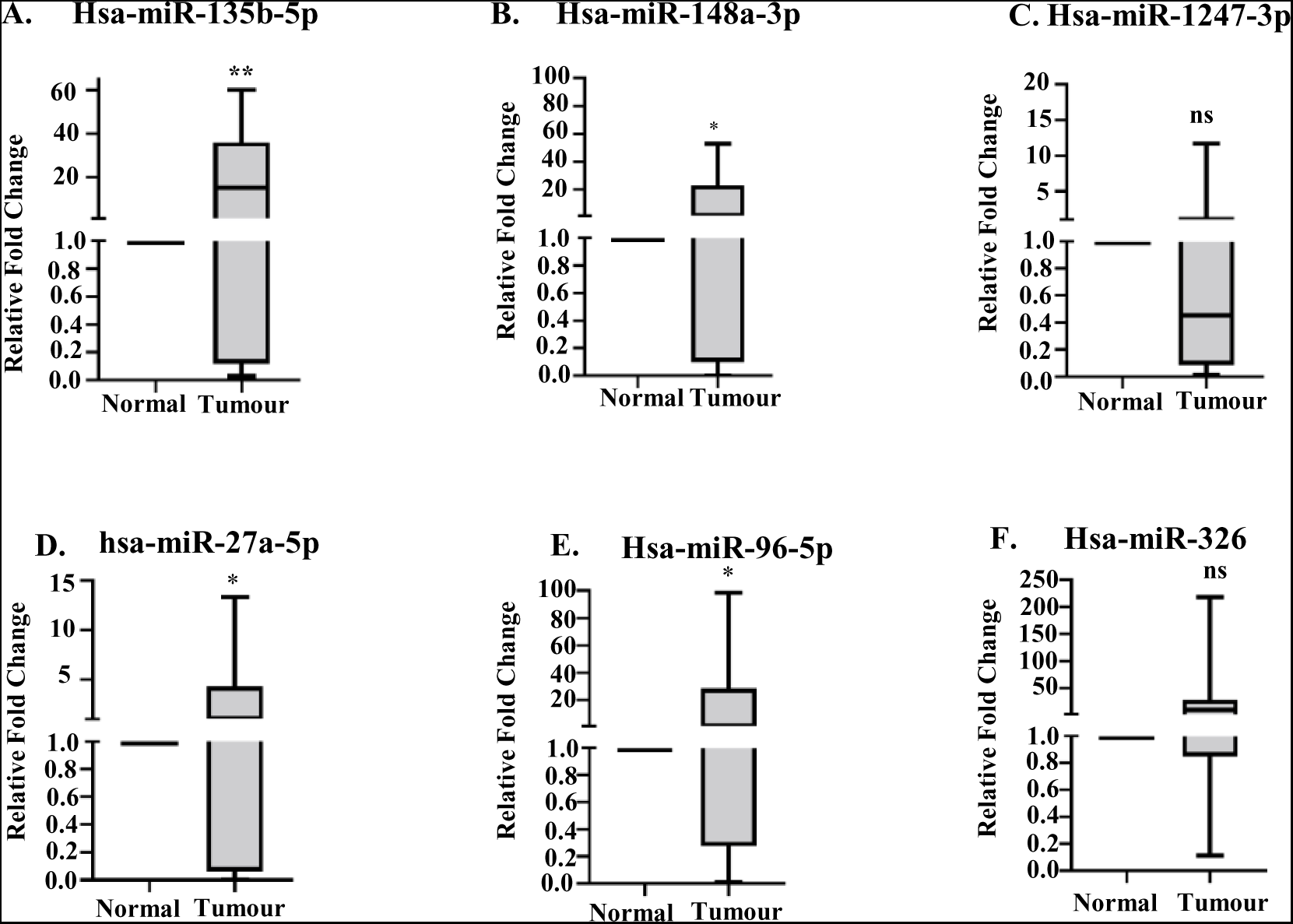
qRT-PCR based differential analysis of young patient validated DEMs in an additional AOCRC validation cohort. Relative expression analysis of **(A)** hsa-miR-135b-5p, **(B)** hsa-miR-148a-3p, **(C)** hsa-miR-1247-3p, **(D)** hsa-miR-27a-5p, **(E)** hsa-miR-96-5p and **(F)** hsa-miR-326 in tumours and normal tissues of 7 aged patients (>55 years old) of our AOCRC validation cohort. RT-qPCR detection of miRNAs was done from 200 ng of isolated cellular RNA. N=3 replicates for each patient tissue (tumour and normal). Normalization was done by U6 snRNA. Data represents Mean ± SD. Statistical significance was calculated by two-tailed Students’ paired t-test. ns: non-significant, *P < 0.05, **P < 0.01.

### Comparison of selected DEMs between young and old tumours

Validation of the selected DEMs in EOCRC and LOCRC validation cohorts were computed on the basis of their expression in tumour tissues as compared to corresponding normal tissues (Figure 4 and 5). It was observed that the degree of significance for the upregulated miRNAs differed between EOCRC and LOCRC patients suggesting different levels of upregulation in young and old tumours (Figure 4 A, B, D, E vs. 5 A, B, D, E). Therefore, to analyse from a different perspective, we compared the delta Ct values of the selected DEMs between tumours of EOCRC and LOCRC cohort patients.

The level of upregulation of hsa-miR-148a-3p, hsa-miR-1247-3p and hsa-miR-27a-5p in young tumours was observed to be significantly more than in old tumours (Figure 6 B, C and D). Hsa-miR-326 was significantly downregulated in young tumours as compared to aged tumours (Figure 6F), consistent with previous observations (Figure 4F and 5F). We therefore selected hsa-miR-148a-3p, hsa-miR-1247-3p, hsa-miR-27a-5p and hsa-miR-326 as being differentially expressed between young and old patients.

**FIGURE 6.**
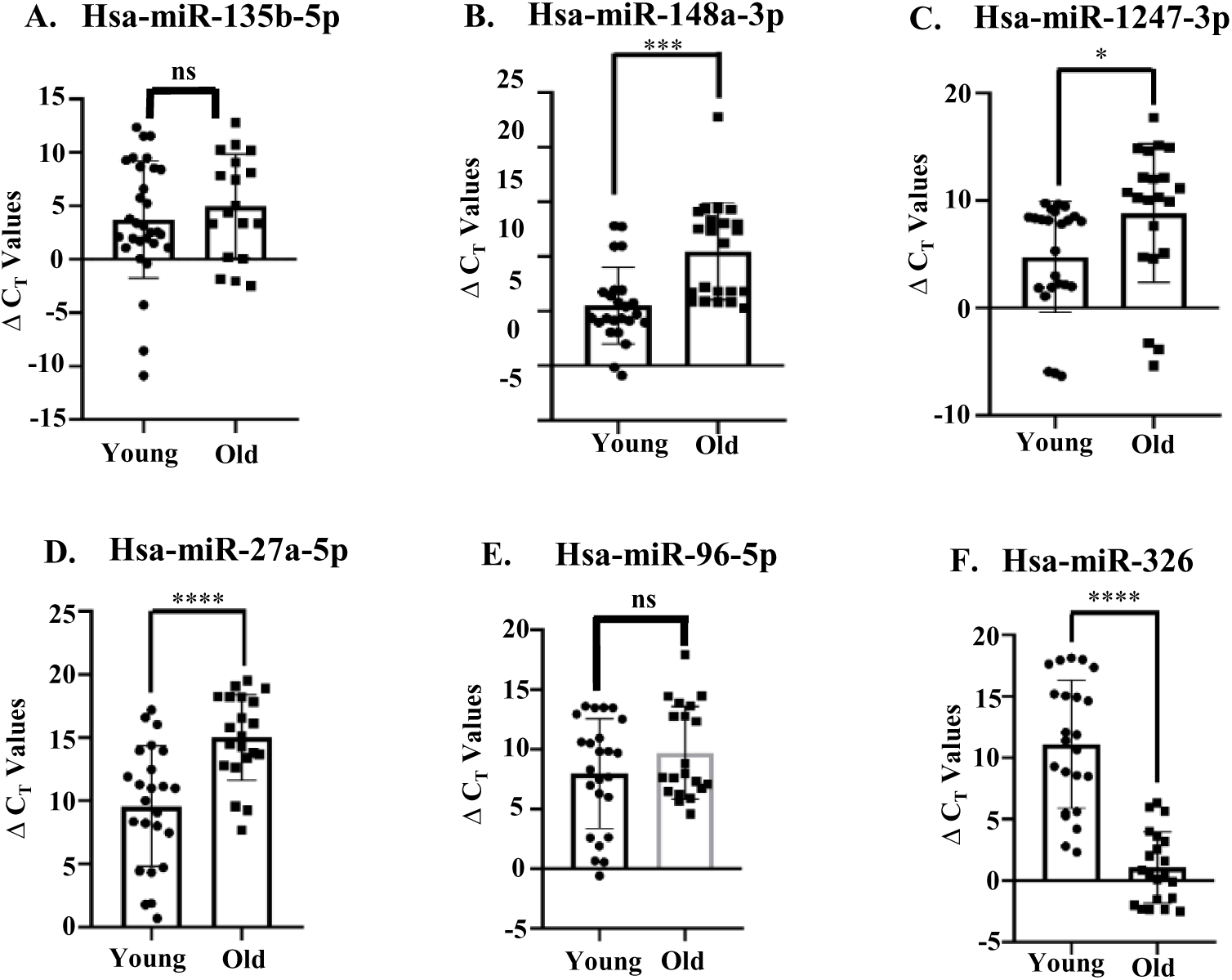
qRT-PCR based differential analysis of validated DEMs in young tumours vs old tumours in additional EOCRC and AOCRC validation cohorts. Delta Ct values of Young tumours (EOCRC validation cohort) were compared with delta Ct values of Aged tumours (AOCRC validation cohort). Relative delta Ct values of **(A)** hsa-miR-135b-5p. **(B)** hsa-miR-148a-3p, **(C)** hsa-miR-1247-3p, **(D)** hsa-miR-27a-5p, **(E)** hsa-miR-96-5p and **(F)** hsa-miR-326 in tumour tissues of 10 young patients and 7 aged patients, n=3 biological replicates for each. Delta Ct= CtmiRNA – CtU6. RT-qPCR detection of miRNAs was done from 200 ng of isolated cellular RNA. Normalization was done by U6 snRNA. Data represents Mean ± SD. Statistical significance was calculated by two-tailed Students’ unpaired t-test. ns: non-significant, *P < 0.05, ***P < 0.001, ****P < 0.0001

### Analysis of DEM targets differentially expressed in TCGA COAD dataset

Experimentally validated targets of hsa-miR-148a-3p, hsa-miR-1247-3p, hsa-miR-27a-5p and hsa-miR-326 were identified through miRNet (https://www.mirnet.ca/) (**17**). Differentially expressed (DE) genes in the TCGA dataset of colorectal adenocarcinomas (COAD) samples with |Log2FC| cutoff of 1.00 and q-value cutoff of 0.01 were identified using GEPIA (http://gepia.cancer-pku.cn/). Target genes of the selected miRNAs were compared by venn-diagram analysis ((https://bioinfogp.cnb.csic.es/tools/venny/) with this DE-gene list to identify DE-target genes altered in colorectal tumour tissue in a direction reciprocal to that of the corresponding miRNA.

Figure 7B depicts the overlap between TCGA-COAD downregulated genes and targets of upregulated miRNAs. 58 genes were common between the two indicating that these targets of the upregulated miRNAs hold physiological significance in colon cancer (Figure 7B). Similarly, we also compared upregulated genes in TCGA COAD dataset with targets of downregulated miRNA hsa-miR-326 (Figure 7C). 61 genes were identified as being common between the two sets indicating their significance in colorectal cancer. Identity of the downregulated 58 targets and upregulated 61 targets have been given in Supplementary Table 3.

**FIGURE 7.**
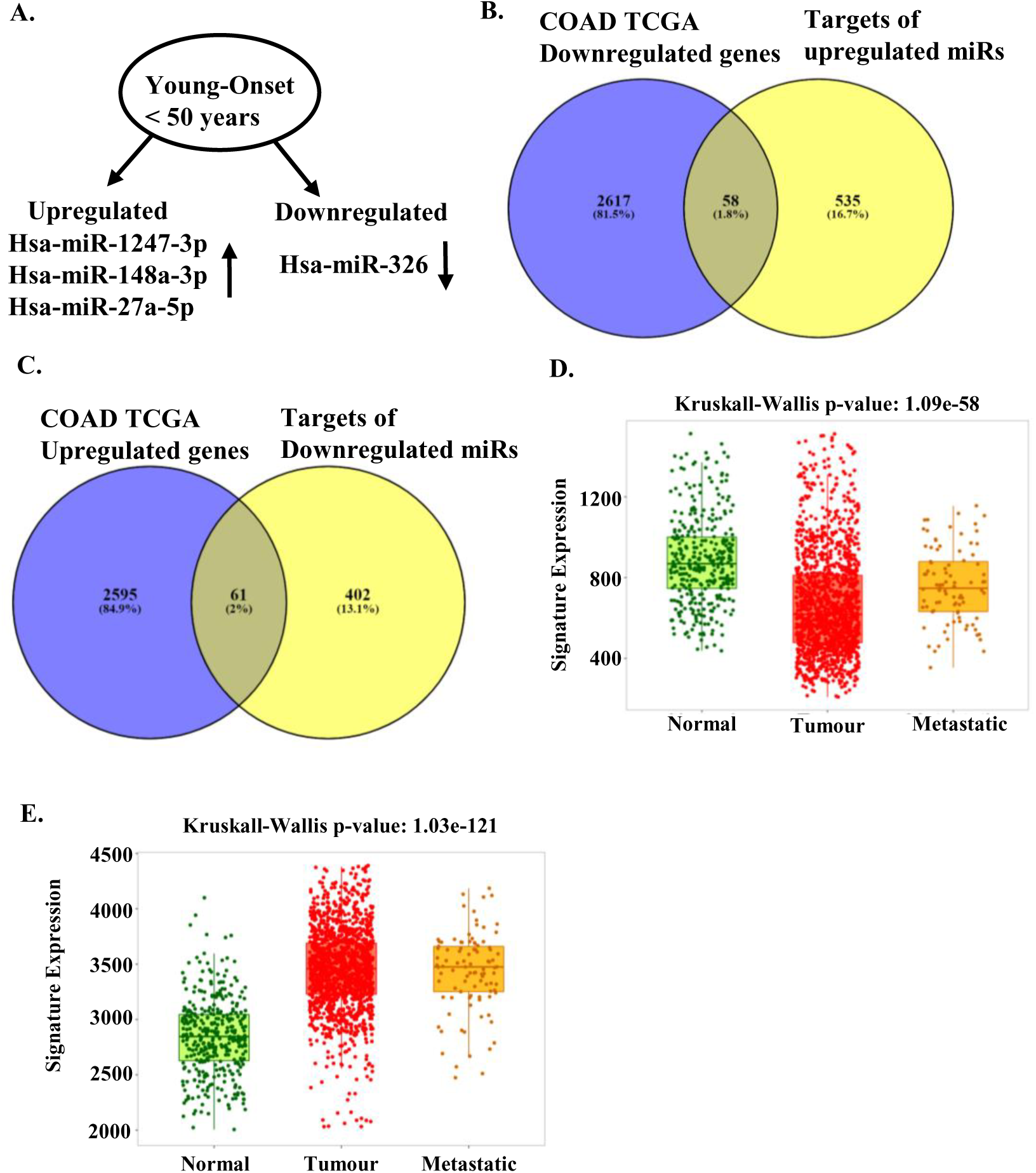
Analysis of upregulated and downregulated DEM target genes in colon cancer development and metastasis. **(A)** Schematic representation of validated upregulated and downregulated DEMs in EOCRC **(B)** Venn diagram comparison of downregulated genes in TCGA dataset of colon adenocarcinoma (COAD) with experimentally verified target genes of upregulated DEMs. 58 target genes were identified as common between the two subsets. **(C)** Venn diagram comparison of upregulated genes in TCGA dataset of colon adenocarcinoma (COAD) with experimentally verified targets of downregulated DEM. 61 target genes were identified as common between the two subsets. **(D)** TNM (Tumour, Normal and Metastasis) plot (https://tnmplot.com/analysis/) of the predicted 58 COAD-downregulated DEM target genes (in B). Differential gene expression analysis of the predicted COAD-downregulated DEM target genes in Tumor, Normal and Metastatic tissues using gene-chip data. Significant downregulation of gene expression was observed between Normal and Tumour and Normal and Metastatic tissues indicating importance of the predicted target genes in tumor development and metastasis. **(E)** TNM (Tumour, Normal and Metastasis) plot (https://tnmplot.com/analysis/) of the predicted 61 COAD-upregulated DEM target genes (in C). Differential gene expression analysis of the predicted COAD-upregulated DEM target genes in Tumor, Normal and Metastatic tissues using gene-chip data. Upregulation of expression of target genes was observed between Normal and Tumour and Normal and Metastatic tissues indicating importance of the upregulation in tumor development and metastasis.

We performed a differential gene expression analysis of these COAD downregulated and upregulated targets in Tumor, Normal and Metastatic colon tissue datasets on TNMplot (https://tnmplot.com/analysis/) (**23**) (Figure 7D and E). Selected targets were downregulated (Figure 7D) and upregulated (Figure 7E) in both tumour and metastatic tissues as compared to normal indicating their role in oncogenesis and in the metastatic spread of disease.

### Pathway enrichment of TCGA-COAD targets of validated DEMs in EOCRC

Gene Ontology and Pathway Enrichment Analysis was performed for the genes common between TCGA COAD datasets and targets of our validated differentially expressed miRNAs to identify pathways deregulated in EOCRC. Enrichment analysis was done using the tool ShinyGO (v.0.75) (**19**) (http://bioinformatics.sdstate.edu/go/) based on the annotations Biological Process (BP), Molecular Function (MF) and Cellular Component (CC) with a FDR cut-off of 0.001 against human as a selected species.

The validated DEM targets for upregulated miRNAs analysed for BP were mainly enriched in Tissue morphogenesis, Morphogenesis of epithelium, Tube development and Epithelium development (Figure 8 A). No enrichment was observed for the annotations Cellular Component and Molecular Function. Validated DEM targets for downregulated miRNAs were enriched the in Biological Process of Protein localization to organelle (Figure 8 B). CC analysis identified the target genes as belonging to ribonucleoprotein complexes (Figure 8 C), the MF category analysis indicated RNA binding proteins (Figure 8 D).

**FIGURE 8.**
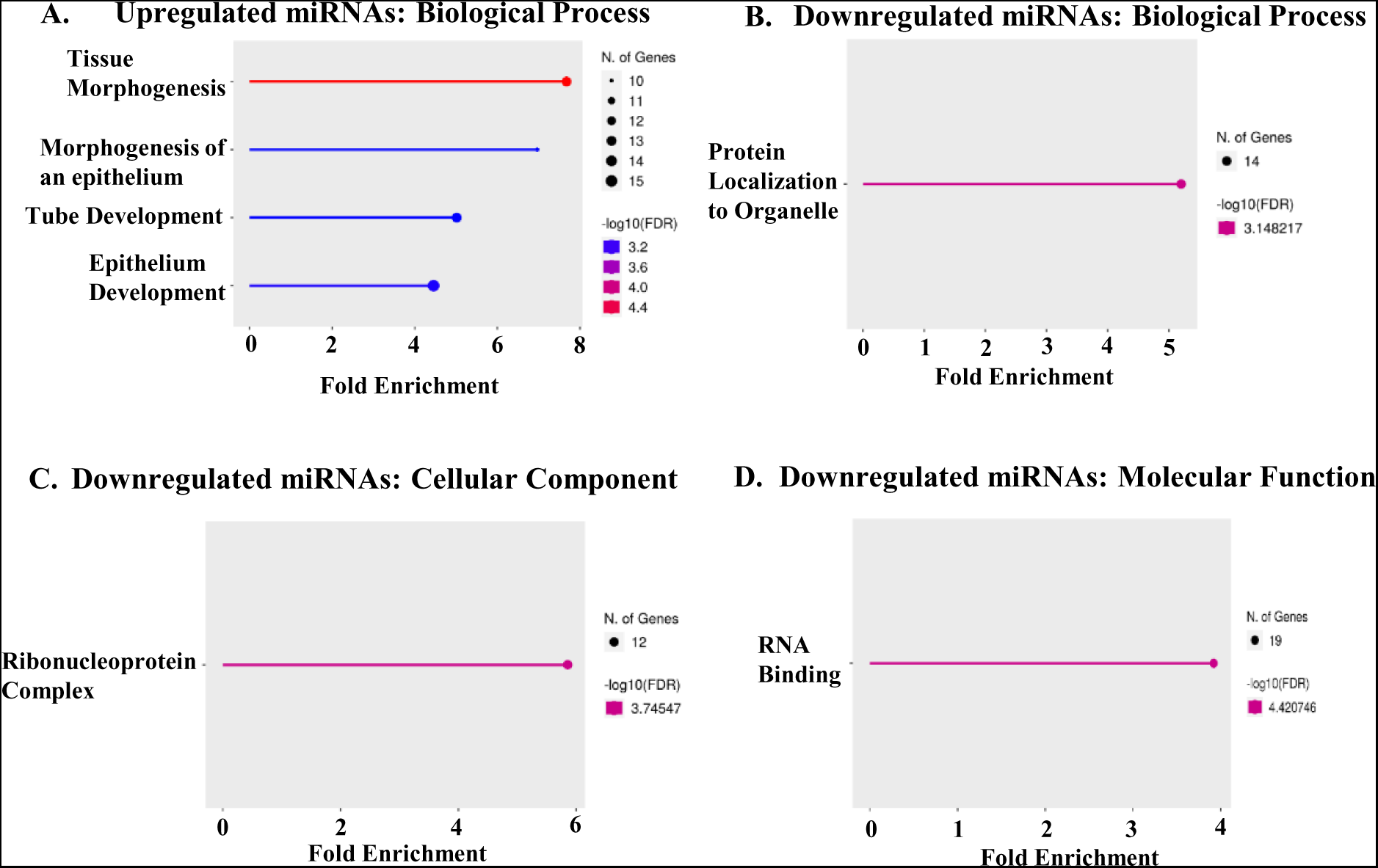
Enrichment analysis of common targets between validated miRs and TCGA COAD database (upregulated and downregulated reciprocally to each other) using the tool ShinyGO (v.0.77) ShinyGO 0.77 (sdstate.edu)) **(A)** represents the gene ontology enrichment chart for the annotation Biological Process (BP) for upregulated miRNAs, **(B)** represents the enrichment chart for BP for downregulated miRNAs **(C)** represents the gene ontology enrichment chart for the annotation Cellular Component and **(D)** represents that for the annotation Molecular Function for targets of validated downregulated miRNA hsa-miR-326. FDR Cut-off was taken as 0.001. Genes have been selected by FDR and sorted by Fold Enrichment. The size of the solid circle indicates the number of targets for each pathway.

## Discussion

The last few decades have seen an increase in the incidence of colorectal cancer among individuals less than 50 years old, also referred to as EOCRC. However not much is known about the aetiology of EOCRC or the reason for its increase in the younger population. Young onset disease was earlier thought to be mainly due to familial causes, however literature indicates that non-familial EOCRC is increasing with clinical and pathological parameters different from classical CRC. miRNAs are short non-coding RNAs that can regulate mRNA levels either positively or negatively and have been shown to be important drivers of disease progression. There is growing interest in exploiting these molecules as either biomarkers for diagnosis, disease prognosis and response to treatment or from a therapeutic perspective as artificial mimics or repressors.

We wanted to perform a molecular analysis of sporadic colorectal tumors in young patients (≤ 50 years old) negative for canonical CRC markers like MSI, nuclear β-catenin and *APC* mutation. To that purpose we screened and collected 5 colorectal tumours with no family history of CRC; 3 belonging to EOCRC (<50years) and 2 belonging to LOCRC (>55 years) and performed small RNA-Seq analysis to identify the unique miRNA signature of each tumor subset. RNA-seq data from these samples were then analysed bioinformatically to identify miRNAs differentially expressed with respect to paired adjacent colonic mucosa (normal) in sample comparisons of Young Normal vs Tumour (EOCRC normal dataset as control group and EOCRC tumour dataset as test group) and Old Normal vs Tumour (LOCRC normal dataset as control group and LOCRC tumour dataset as test group). Venn diagram comparison of DEMs in ‘Young Normal vs Tumour’ (EOCRC) and ‘Old Normal vs tumour’ (LOCRC) highlighted miRNAs specific to each subtype and also those common to both subtypes. 23 miRNAs were differentially expressed specifically in young patients, 11 miRNAs were differentially expressed specific to aged patients and 5 miRNAs were found to be differentially expressed in both. The 5 miRNAs (hsa-miR-129-5p, hsa-miR-9-5p, hsa-miR-1-3p, hsa-miR-145-5p and hsa-miR-133a-3p) common to both EOCRC and LOCRC have been previously reported as downregulated in CRC (**24**, **25**, **26**). Top 10 EOCRC DEMs were first validated in TCGA colorectal adenocarcinoma (COAD) dataset using miRNome (<u>CancerMIRNome (jialab-ucr.org)</u>) and differentially expressed miRNAs upregulated or downregulated in COAD in the same direction as our NGS analysis were then detected in additional EOCRC and LOCRC cohorts by RT-qPCR to select significantly upregulated and downregulated miRNAs specific to EOCRC. Upregulated DEMs included hsa-miR-1247-3p, hsa-miR-148a-3p and hsa-miR-27a-5p. Hsa-miR-326 was significantly downregulated in EOCRC.

Experimentally validated targets of the selected miRNAs were compared with genes differentially expressed (DE) in TCGA COAD samples to identify predicted DE-target genes altered in colorectal tumour tissue in a direction reciprocal to that of miRNAs. We identified 58 downregulated (miR upregulated) and 61 upregulated (miR downregulated) DE target genes. These predicted DE-targets were analysed by TNMplot software to confirm upregulation/ downregulation in Tumor, Normal and Metastatic colon tissue datasets. GO pathway analysis of predicted DEM targets known to be differentially expressed in colon adenocarcinoma (TCGA-COAD) was conducted using SHINYGO online software with a FDR cut-off of 0.001. DE target genes of upregulated miRNAs were enriched in pathways of tissue morphogenesis, morphogenesis of epithelium, tube development and epithelium development. Downregulated miRNA DE target genes were enriched in the biological process of protein localization to organelle, belonged to the cellular compartment of ribonucleoprotein complexes with molecular function of RNA binding.

Predicted downregulated targets include negative regulators of the WNT pathway-SFRP1 and PRICKLE1 (**27**, **28**), the tumor suppressor IGFBP5 which shows inverse correlation with IGF1R phosphorylation (**29**, **30**), the GTPase activating protein STARD13 which has roles in cell cycle inhibition (**31**), apoptosis (**32**) and as a regulator of CDC42 which plays a major role in epithelial tissue formation and homeostasis, invasion, extracellular matrix degradation and mitotic progression (**33**). Also predicted to be downregulated is the epithelial intercellular junction protein ADD1 involved in remodelling of epithelial junctions and restoration of epithelial barrier (**34**, **35**). This has important physiological significance as downregulation of epithelial gene expression signature and the dissolution of epithelial intercellular junctions are key events in epithelial-mesenchymal transition (**36**).

Predicted upregulated targets in the BP annotation “Protein localization to organelle” include various molecules whose expression are known to correlate with the parameters of disease aggressiveness like tumor invasion, liver metastasis, disease recurrence and poor prognosis. These include the transcription factor XBP1 required for MHC Class II gene regulation (**37**, **38**) angiogenesis (**39**, **40**) and ER stress response (**38**, **41**), BMP7 which is a secreted signalling molecule belonging to the TGF-β superfamily (**42**), the transcription factor E2F1 that regulates cell cycle progression (**43**), the ribosomal protein RPL28 that is a predictor of survival in metastatic CRC patients (**44**), the CRC prognostic marker NPM1 that activates AKT signalling (**45**) and the

RHO GTPase activator ECT2 that plays important driver roles in CRC oncogenesis (**46**). The CC annotation shows upregulation of targets belonging to “Ribonucleoprotein complexes” like eukaryotic ribosome biogenesis protein hMRT4/ MRTO4 (**47**), the pre-mRNA splicing factor EFTUD2 that is a marker of chronic intestinal inflammation and tumorigenesis as it modulates TLR4 signaling and NF-κB activation in macrophages (**48**), the ER chaperone protein CALR which is a key regulator of Ca2+ homeostasis and integrin-dependent signalling (**49**) and RPL28

(**44**). Upregulated “RNA binding” proteins in the annotation of Molecular Function include MRTO4 (**47**), RPL28 (**44**), proteins involved in the metabolic shift from oxidative phosphorylation to aerobic glycolysis like Pyruvate Kinase M1/2 (PKM) and Heterogeneous Nuclear Ribonucleoprotein A1 (HNRNPA1) (**50**), the oncogene and colon carcinoma prognostic marker glutamate-rich WD40 repeat containing 1 (GRWD1) (**51**), Elongation Factor Tu GTP Binding Domain containing 2 (EFTUD2) (**48**) and the prognostic and metastatic marker Fatty Acid Synthase (FASN) (**52**). All these target genes have important roles in the development and progression of colorectal cancer.

In spite of the bulk of previously conducted studies on CRC gene expression and miRNAs (**53, 54, 55. 56, 57, 58, 10, 11**), there are significant gaps in our understanding of EOCRC and how it differs from LOCRC. Most of the published reports with genome-wide RNA-sequencing (**54**, **11**) concentrate on a cohort comprised exclusively of EOCRC patients. The inclusion of sporadic LOCRC is essential in the initial discovery cohort as it is not possible to identify markers specifically deregulated in early onset disease (significantly upregulated/ downregulated with respect to LOCRC) without comparison with LOCRC tissues. The diagnosis of CRC before the age of 50 always raises the suspect of a genetic cancer predisposition syndrome (Lynch Syndrome and Familial Adenomatous Polyposis). So it is important to screen the EOCRC/ LOCRC cohort for known canonical markers as their presence creates a hyper-mutable and pro-oncogenic phenotype. Liu et al. performed a genome wide miRNA and transcriptome profiling but the tumours included in their study cohorts were not assessed for the canonical CRC markers like MSI or activation of the Wnt pathway or *APC* mutations (**58**). Another bottle-neck of big-data transcriptomic studies is that they very rarely include paired adjacent colonic mucosa as normal samples in their analysis. Ideally tumour samples should be paired with corresponding normal samples to avoid biological differences between individuals.

Our study is the first attempt to identify differentially expressed miRNAs specific to EOCRC in the Indian population. We have attempted to remove the inconsistencies of previous studies by the inclusion of a LOCRC cohort both in the discovery cohort and also in the validation cohort with paired adjacent colonic mucosa corresponding to each tumor analyzed in the study. Additionally we have also screened our cohorts for MSI, nuclear β-catenin and *APC* mutations to collect tumors negative for these known CRC canonical markers. Since a miRNA can target many mRNAs we screened the validated targets of our EOCRC validated DEMs to identify those target genes (58 downregulated and 61 upregulated) known to be differentially expressed in colorectal adenocarcinoma datasets. In future, these target genes need to be validated in EOCRC and LOCRC cohorts for identification of potential pathways responsible for the early onset of colorectal cancer.

### Data Availability Statement

The data discussed in this publication have been deposited in NCBI’s Gene Expression Omnibus (**60**) and are accessible through GEO Series accession number GSE236991 (https://www.ncbi.nlm.nih.gov/geo/query/acc.cgi?acc=GSE236991). Validation data supporting the findings of this study are available from the corresponding authors upon request.

## Acknowledgements

The work received support from a Science and Engineering Research Board-Start Up Research Grant (SERB-SRG) (SRG/2019/000639) from Science and Engineering Research Board, Govt. of India. SuM acknowledges the support from Indian Council of Medical Research, Govt. of India for her Fellowship support. We thank Dr. Rittwika Bhattacharya, Senior Scientist, Department of Molecular Biology, Netaji Subhas Chandra Bose Cancer Research Institute, Kolkata, for instrumental support. We thank Mr. Sourav K. Nandi and Ms. Minakshi Sen for providing support in methodology and instrument use. We acknowledge methodological support from Dr. Sumita Sengupta, Assistant Professor, Department of Biophysics, Molecular Biology and Bioinformatics, University of Calcutta. We acknowledge all patronage from late Dr Ashis Mukhopadhyay, former Director, Netaji Subhas Chandra Bose Cancer Research Institute, Kolkata. We acknowledge Dr. Suvendra N. Bhattacharyya and his student, Mr. Diptankar Bandyopadhyay, RNA Biology Research Laboratory, Molecular Genetics Division, CSIR-Indian Institute of Chemical Biology, Kolkata, for allowing us access to the NanoDrop facilities for nucleic acid quantification.

## Author Contributions

SB and SG conceived the idea, designed the experiments and analyzed the data. SuM, VM and DB performed the experiments. SD and BB were the clinicians associated with the study. NB, AD and KC were the histopathologists associated with the study. BD provided technical help and support for the study. SM provided support with project administration and resources. SB and SG wrote the manuscript. SuM also contributed in analyzing the data and writing the manuscript.

