## Supplementary Data for "Differential expression of miRNAs between Young-Onset and Late-Onset Indian colorectal carcinoma patients"

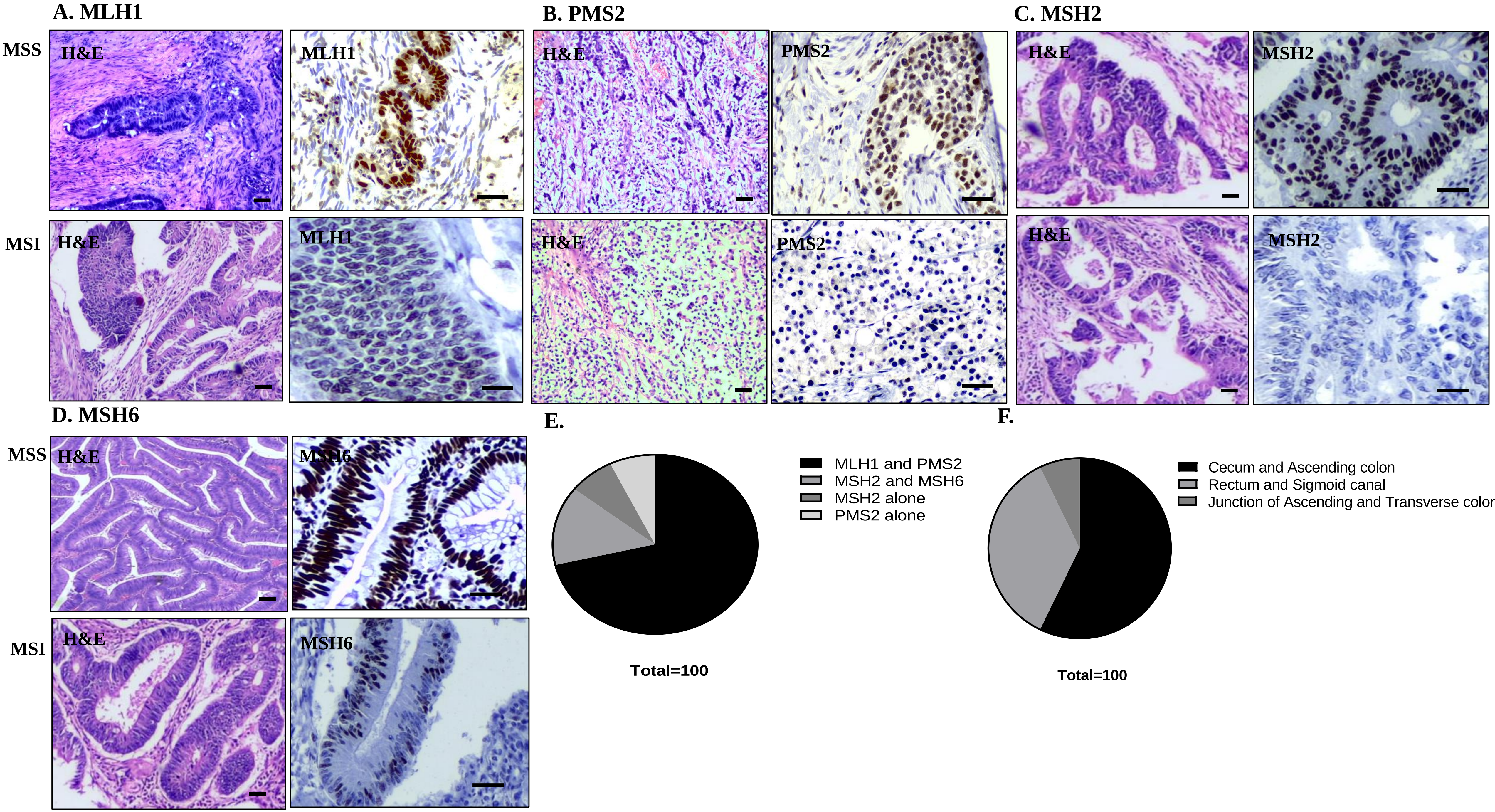


**FIGURE S1 IHC based detection of Microsatellite instability markers (MLH1, PMS2, MSH2 and MSH6) in MSI and MSS colorectal tumour tissues**

**(A – D)** Representative micrograph images showing depleted MMR protein in MSI and MSS colorectal tumour tissues. Depicted Haematoxylin & Eosin images are in 10x magnification and IHC images are in 20x magnification. Scale bar represents 100 µm. **E.** Percentage-wise distribution of depleted MMR protein in MSI colorectal tumour tissues. **F.** Percentage-wise distribution of location of dMMR tumours.

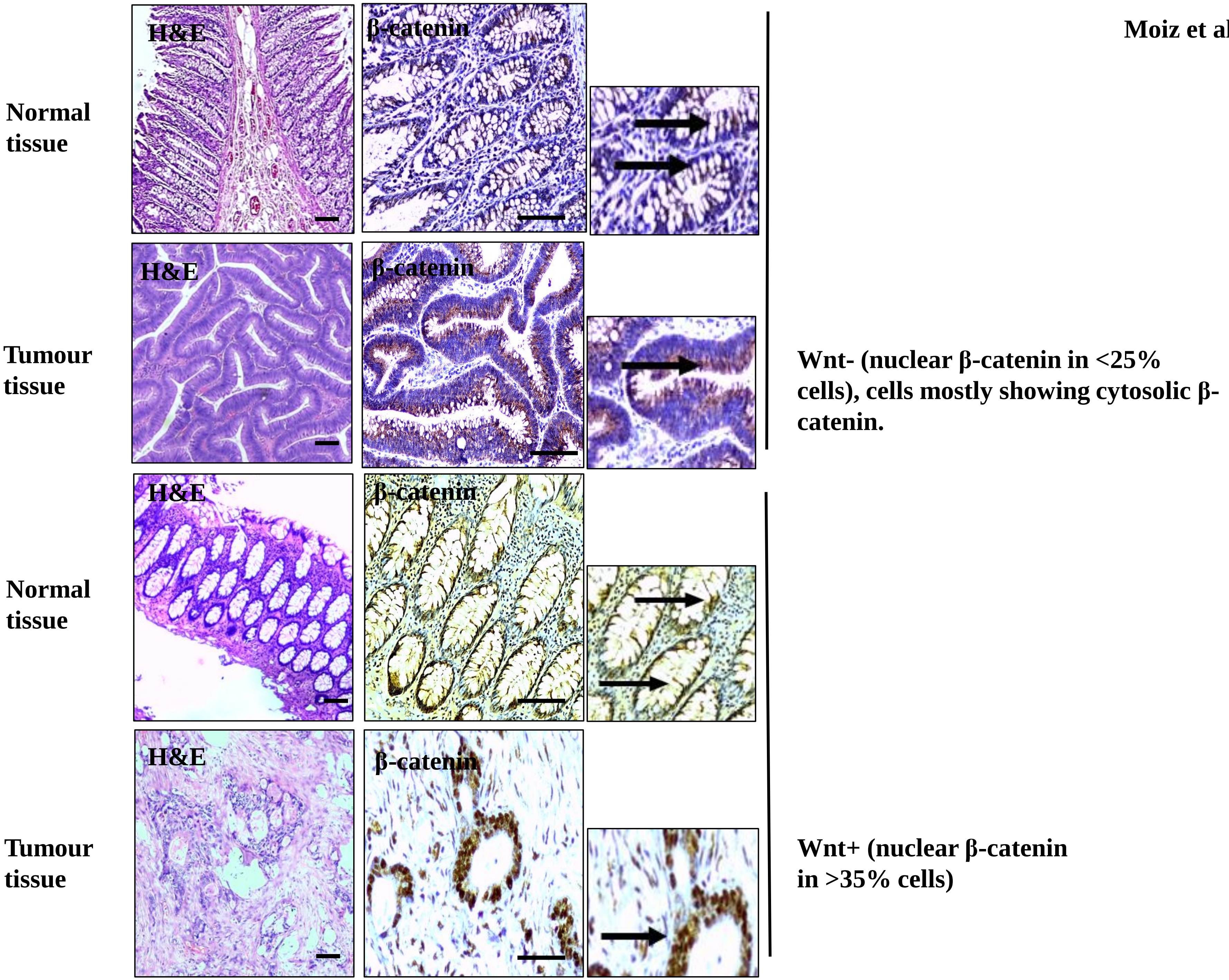


**FIGURE S2 IHC based detection of β-catenin in normal and tumour tissues**

Representative micrograph images showing IHC based detection of β-catenin in normal and tumour tissues. Normal tissues show membrane localization of β-catenin (inset arrows). Wnt- tumour tissues show mainly cytoplasmic localization of β-catenin (inset arrows). Wnt+ tissues show nuclear β-catenin in >35% cells. Areas marked by black squares have been zoomed into with arrows showing membrane, cytosolic or nuclear localization of β-catenin. Depicted Haematoxylin & Eosin images are in 10x magnification and IHC images are in 20x magnification. Scale bar represents 100 µm.

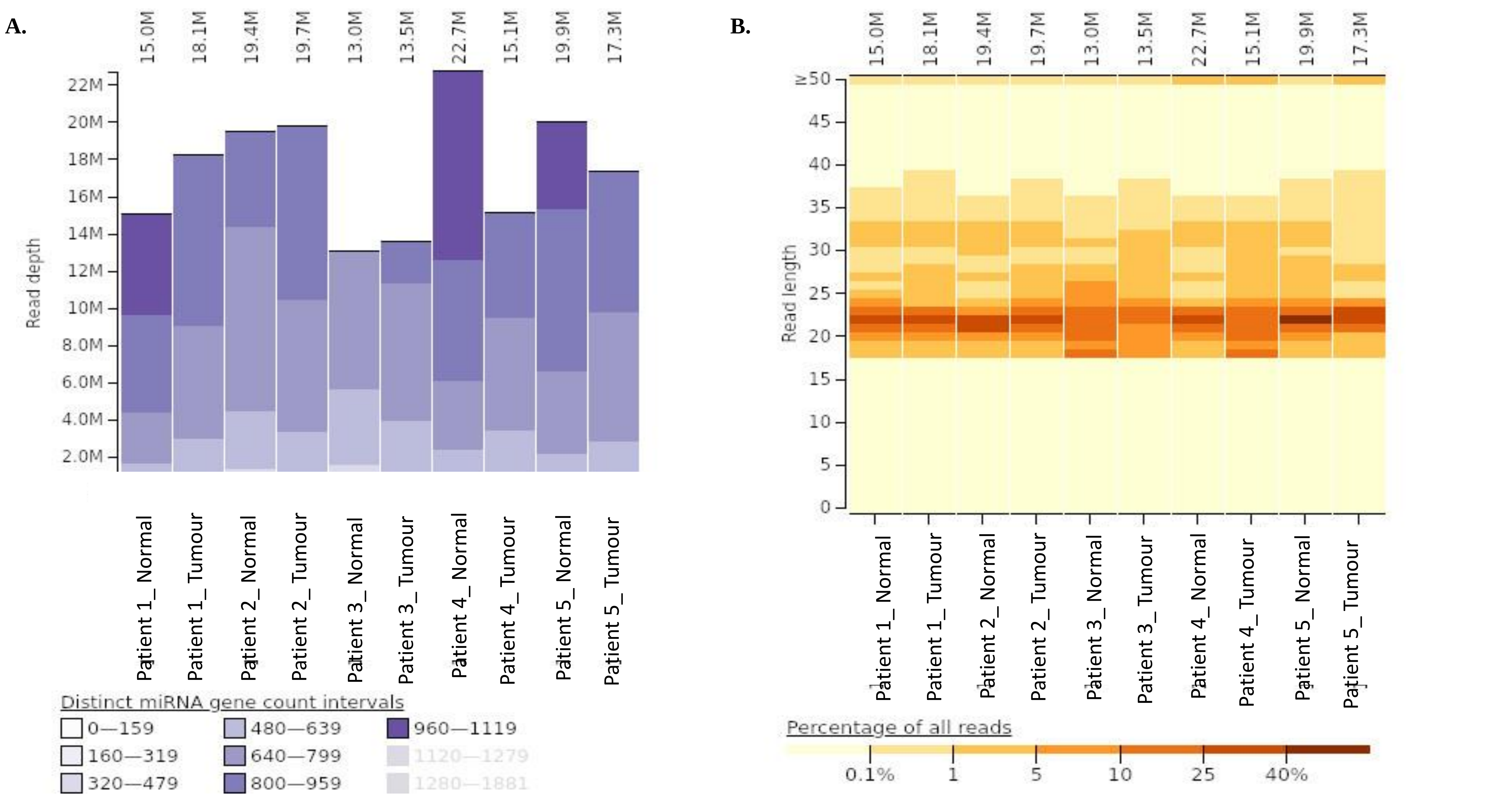


**Figure S3 Read Depth Coverage and Sequence Length Distribution Plots**

**(A)** Read depth coverge was estimated using miRTrace v.1.0.1 [59]. Read depth coverage plot showed the read depth (no. Of reads in million) at the y-axis and samples at the x-axis. **(B)** Sequence length distribution was estimated using the miRTrace. Bar diagram was generated using the cleaned sequences of samples. Over 40% sequences in each sample were found to have length distribuion from 18 nt. (minimum trimmed length) to 23 nt.

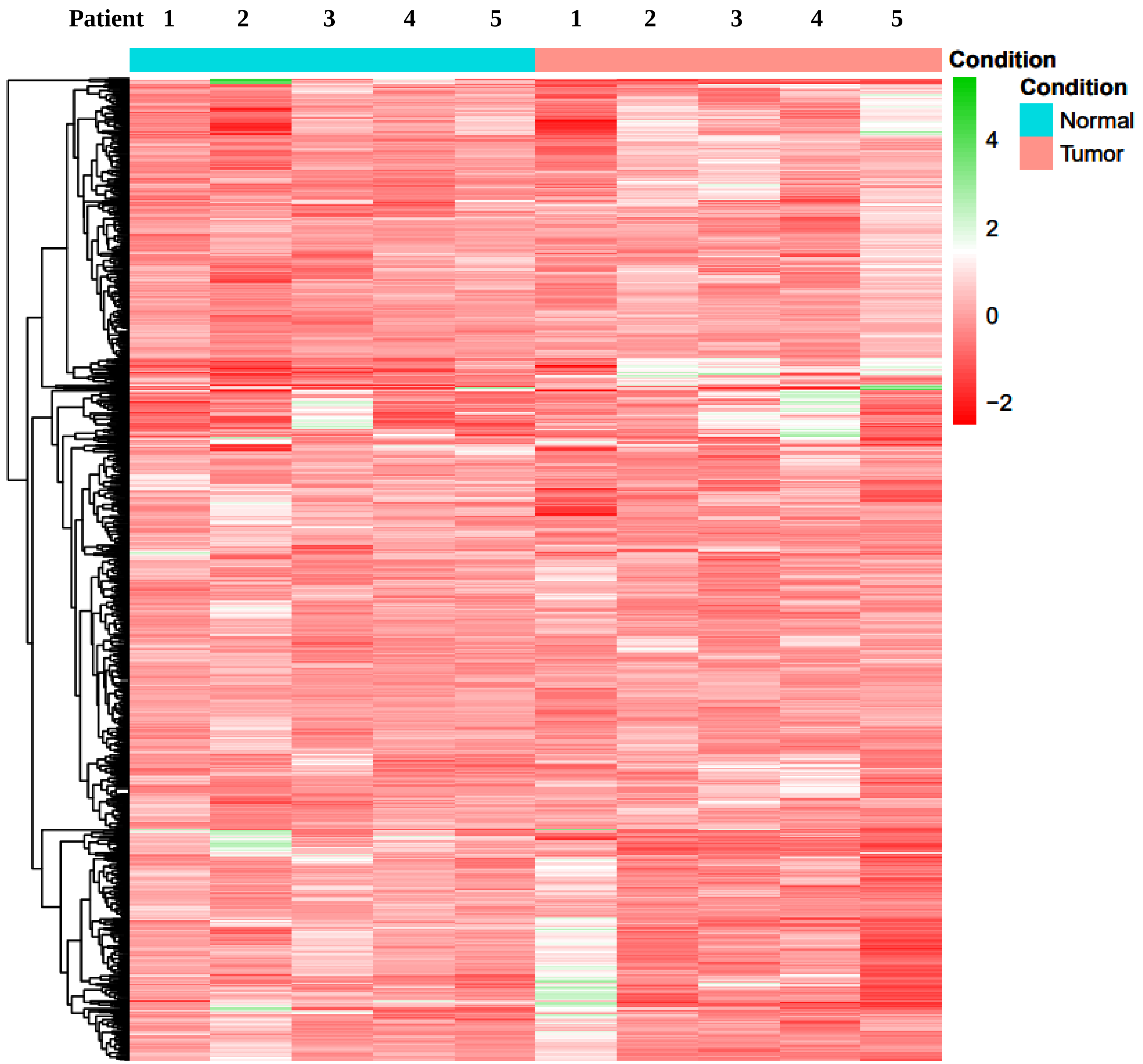


**Figure S4 Heatmap of all miRNAs sorted by variance in decreasing order, found across 10 samples (5 normal and 5 tumour)**

The heatmap was generated by using rlog-normalized count data of each sample. Z-scores for each miRNA per sample are indicated in red and green colour scale shown on the right, for expression levels below and above mean expressions across sample for that given miRNA, respectively. The condition legend denotes the sample types, where cyan blue is assigned to Normal samples and pink is assigned to Tumor samples.

**Supplementary Table 1:**

**Detailed information on the sequencing miRNA library of CRC tissues: tumor and paired normal tissues.**

| Sample | Total Reads | Cleared Reads | % Cleared Reads | Total Sequences | Mapped Sequences | % Mapped Sequences |
|---|---|---|---|---|---|---|
| 10_normal | 15609568 | 14952600 | 95.79% | 14747826 | 5359822 | 36.343% |
| 10_tumour | 20282568 | 18134722 | 89.41% | 17861428 | 5506910 | 30.831% |
| 13_normal | 20016594 | 19448545 | 97.16% | 19178794 | 6863344 | 35.786% |
| 13_tumour | 21324902 | 19747475 | 92.60% | 19457532 | 6064884 | 31.170% |
| 14_normal | 15102654 | 12986956 | 85.99% | 12784955 | 1038689 | 8.124% |
| 14_tumour | 15628536 | 13482241 | 86.27% | 13274675 | 2170200 | 16.348% |
| 15_normal | 23309080 | 22718559 | 97.47% | 22409421 | 8376465 | 37.379% |
| 15_tumour | 16669194 | 15058561 | 90.34% | 14830961 | 1451003 | 9.784% |
| 20_normal | 20589592 | 19923833 | 96.77% | 19631591 | 7282762 | 37.097% |
| 20_tumour | 17753430 | 17296336 | 97.43% | 17045393 | 6762325 | 39.672% |

**Supplementary Table 2:**

**List of significant differentially expressed miRNAs identified in multiple sample comparisons of miRNA sequencing results of young (<50 years old) and aged (>55 years old) colorectal cancer apatient tumours (DEM Cut Off: p-value <0.05, Log Fold Change: >2, <-2)**

| Comparison (Sample_vs_Control) | miRNA identity | $Log_2FC$ | P-value | Up ↑ / Down ↓ |
|---|---|---|---|---|
| All Tumour vs All Normal | hsa-miR-9-5p | -3.08529936 | 1.73E-05 | Down ↓ |
| All Tumour vs All Normal | hsa-miR-129-5p | -3.46090279 | 3.41E-05 | Down ↓ |
| All Tumour vs All Normal | hsa-miR-1-3p | -4.082048456 | 6.43E-05 | Down ↓ |
| All Tumour vs All Normal | hsa-miR-133a-3p | -4.139735223 | 0.000118847 | Down ↓ |
| All Tumour vs All Normal | hsa-miR-145-5p | -2.927381878 | 0.000286046 | Down↓ |
| Young Tumour vs Young Normal | hsa-miR-1247-3p | 3.70881538 | 0.00174089 | Up↑ |
| Young Tumour vs Young Normal | hsa-miR-27a-5p | 2.61304708 | 0.00901648 | Up↑ |
| Young Tumour vs Young Normal | hsa-miR-96-5p | 2.80991462 | 0.02211184 | Up↑ |
| Young Tumour vs Young Normal | hsa-miR-148a-3p | 2.49604406 | 0.02365696 | Up↑ |
| Young Tumour vs Young Normal | hsa-miR-135b-5p | 3.41150654 | 0.02799316 | Up↑ |
| Young Tumour vs Young Normal | hsa-miR-378e | -2.61154969 | 0.00366325 | Down↓ |
| Young Tumour vs Young Normal | hsa-miR-326 | -2.63728355 | 0.01712207 | Down↓ |

| | | | | |
|---|---|---|---|---|
| **Young Tumour vs Young Normal** | **hsa-miR-378c** | **-2.49476989** | **0.01533526** | **Down↓** |
| **Young Tumour vs Young Normal** | **hsa-miR-378a-5p** | **-2.48712807** | **0.00620528** | **Down↓** |
| **Young Tumour vs Young Normal** | **hsa-miR-378d** | **-2.41909916** | **0.01914212** | **Down↓** |
| **Young Tumour vs Young Normal** | **hsa-miR-363-3p** | **-2.40777269** | **0.04421748** | **Down↓** |
| **Young Tumour vs Young Normal** | **hsa-miR-143-3p** | **-2.29934116** | **0.01331962** | **Down↓** |
| **Young Tumour vs Young Normal** | **hsa-miR-504-5p** | **-2.28494574** | **0.01904966** | **Down↓** |
| **Young Tumour vs Young Normal** | **hsa-miR-139-5p** | **-2.27780576** | **0.0240824** | **Down↓** |
| **Young Tumour vs Young Normal** | **hsa-miR-139-3p** | **-2.1809868** | **0.00568353** | **Down↓** |
| **Young Tumour vs Young Normal** | **hsa-miR-9-3p** | **-2.15316894** | **0.00770244** | **Down↓** |
| **Young Tumour vs Young Normal** | **hsa-miR-490-5p** | **-2.14003278** | **0.01709929** | **Down↓** |
| **Young Tumour vs Young Normal** | **hsa-miR-133a-5p** | **-2.13259517** | **0.00324305** | **Down↓** |
| **Young Tumour vs Young Normal** | **hsa-miR-490-3p** | **-2.11550509** | **0.01130573** | **Down↓** |
| **Young Tumour vs Young Normal** | **hsa-miR-887-3p** | **-2.10734608** | **0.00627189** | **Down↓** |
| **Young Tumour vs Young Normal** | **hsa-miR-30a-5p** | **-2.0469516** | **0.00769662** | **Down↓** |
| **Young Tumour vs Young Normal** | **hsa-miR-133b** | **-2.01434615** | **0.00292072** | **Down↓** |

| | | | | |
|---|---|---|---|---|
| **Young Tumour vs Young Normal** | **hsa-miR-143-5p** | **-2.0070137** | **0.02078392** | **Down↓** |
| **Old Tumour vs Old Normal** | **hsa-miR-455-3p** | **2.01483885** | **0.008717** | **Up↑** |
| **Old Tumour vs Old Normal** | **hsa-miR-31-3p** | **3.06621503** | **0.01037511** | **Up↑** |
| **Old Tumour vs Old Normal** | **hsa-miR-204-5p** | **-4.35495512** | **0.01160546** | **Down↓** |
| **Old Tumour vs Old Normal** | **hsa-miR-1247-5p** | **2.63591607** | **0.02448825** | **Up↑** |
| **Old Tumour vs Old Normal** | **hsa-miR-10524-5p** | **2.07155472** | **0.02805908** | **Up↑** |
| **Old Tumour vs Old Normal** | **hsa-miR-549a-5p** | **3.15057457** | **0.03220999** | **Up↑** |
| **Old Tumour vs Old Normal** | **hsa-miR-934** | **2.11438475** | **0.03480386** | **Up↑** |
| **Old Tumour vs Old Normal** | **hsa-miR-135b-3p** | **2.6736609** | **0.03507436** | **Up↑** |
| **Old Tumour vs Old Normal** | **hsa-miR-10396b-3p** | **2.00250344** | **0.03772035** | **Up↑** |
| **Old Tumour vs Old Normal** | **hsa-miR-4485-3p** | **2.92834074** | **0.04423556** | **Up↑** |
| **Old Tumour vs Old Normal** | **hsa-miR-31-5p** | **4.86227719** | **0.04948822** | **Up↑** |

**Supplementary Table 3:**

**Targets of validated significantly upregulated/ downregulated miRNAs in EOCRC that are differentially expressed in colorectal cancer TCGA datasets (TCGA COAD).**

| TARGET miR UP/ DOWN | TCGA COAD DOWN/UP | SELECTED GENES |
|---|---|---|
| Target miR UP<br>Hsa-miR-1247-3p, hsa-miR-148a-3p, hsa-miR-27a-5p | TCGA COAD DOWN | WNT2B, CPM, DNAJB4, ADD1, BCL2, RBM38, MXI1, ZDBF2, GK5, UNKL, RILPL1, SCD5, SCARF1, STX2, ETFDH, MDM4, GPR183, PBXIP1, PRICKLE1, TNRC6A, ZNF793, KIAA0513, RAB12, TPCN2, VSTM4<br>PLEKHG2, S1PR1, TUBB2A, QKI, IGFBP5, STARD13, NEURL4, SNAP25, PRNP, C3, LIX1L, COLEC12, PBX1, TXNIP, CRISPLD2, PHLDA3, KIAA1614, KIAA0408, RASSF8, ADARB1, NPR1, ZMAT1, BRSK1, RAB34, SSBP2, GAS1, NT5DC3, GLP2R, BMP3, SFRP1, ITGA5, PHLDB2, NPTX1 |
| Target miR DOWN<br>Hsa-miR-326 | TCGA COAD UP | SDC1, IHH, BIRC5, EPHB3, ECT2, BMP7, KPNA2, RBM47, ITGA2, AP1S1, CCND1, E2F1, TMEM33, CALR, RAB3IP, MZT1, MTHFD2, SMIM15, NOB1, ALYREF, CSE1L, NPM1, FASN, PLA2G4F, EIF1AX, PKM, BCL2L1, CNNM4, RPL36, ACTG1, POFUT1, RSL1D1, HMGA2, P4HB, E2F2, ACLY, ULBP3, CYB561D2, HMGCS1, SF3B3, MRTO4, NCLN, GLO1, INSIG1, CENPL, ZBTB33, EFTUD2, RPL27A, EMC10, PPM1G, XBP1, NCL, HNRNPA1, GRWD1, BCL11B, CD276, RPL28, MOCS3, ABCC6, GEMIN6, ACVR1B |
